# Identification and characterization of the delta-12 fatty acid desaturase from *Euglena gracilis*

**DOI:** 10.1101/2025.09.11.675727

**Authors:** Raj Kumar Thapa, Bijaya Kumar Uprety, RJ Neil Emery, Scott C Farrow

## Abstract

Fatty acid desaturase 12 (FAD12) is a key enzyme in fatty acid biosynthesis, responsible for converting oleic acid to linoleic acid through desaturase activity. *Euglena gracilis* (Euglena) is an emerging platform for the industrial production of various metabolites, including lipids. However, a comprehensive understanding of Euglena’s fatty acid biosynthesis pathways remains incomplete, posing a significant barrier to the commercialization of Euglena bioproducts. To address this gap, we employed a bioinformatics approach to identify a *Euglena gracilis* FAD12 (*Eg* FAD12). We analyzed the evolutionary relationship of *Eg* FAD12 with its homologs from other organisms and revealed that the three canonical histidine box motifs are conserved among FAD12s. To characterize *Eg*FAD12, we cloned it into the pEAQ-hyperstrans vector and overexpressed it in *Nicotiana benthamiana* to take advantage of its endogenous fatty acid pool, which could act as substrates. The heterologous expression of FAD12 in *N. benthamiana* led to an increased linoleic acid content, demonstrating the suspected desaturase activity. To further confirm the function of *Eg* FAD12, we performed CRISPR-Cas9-mediated knockout of *Eg* FAD12 in Euglena, which resulted in a drastic reduction in linoleic acid (C18:2) without compromising biomass yield or lipid content. This work advances our understanding of fatty acid biosynthesis in Euglena and will aid in its adoption as a platform for producing customized lipids.

## Introduction

Microorganisms like bacteria, yeasts, fungi, and microalgae produce microbial lipids. They are like traditional plant and animal lipids, but offer several benefits when considering them as a lipid source. This includes i) reduced land and water need, ii) efficient labour, iii) climate change resiliency, iv) faster production cycles, and v) scalability. Fatty acids (FA) are integral components of lipids and play a substantial role in defining their physical and chemical properties. This affects the industrial use of such lipids (Uprety et al., 2022). Therefore, developing methods to modify the FA composition of microbial lipids is vital, allowing the customization of lipids for specific industrial purposes. This strategy can enhance the applications of microbial lipids, making them more economically viable and attractive for commercial use (Ochsenreither et al., 2016).

*Euglena gracilis* (Euglena) is an adaptable microalgae capable of growing phototrophically, heterotrophically, or mixotrophically on a variety of substrates and in harsh environments (Farjallah et al., 2024; Kuhne et al., 2023; Lewis & Guéguen, 2020). Euglena can produce a wide range of commercial bioproducts such as amino acids, pro(vitamins), lipids, and the immunogenic glycopolymer paramylon (Rodríguez-Zavala et al., 2010; Uprety et al., 2022). This makes Euglena an attractive biological platform for food, pharmaceutical, and fuel industries. Besides this, it also has the potential to be used as a biological agent to treat wastewater for sustainable management (Mahapatra et al., 2013) or heavy metal contaminated sites (Nguyen et al., 2023). There are studies demonstrating its medicinal properties, such as wound healing (Ko et al., 2023), next-generation prebiotics (Dai et al., 2022), and a booster of the immune system by activating natural killer cells (Park et al., 2023). In addition, there is growing interest in its use as a producer of sustainable biofuel (Chen et al., 2022; Kim et al., 2023).

*Euglena* can accumulate lipids comprising up to 37% of its dry weight, with a FA profile that varies considerably depending on environmental growth conditions (Shibata et al., 2018; Wang et al., 2018). Among these lipids, oleic acid, linoleic acid, and stearic acid are of particular interest due to their functional applications in biofuels, cosmetics, and nutritional products (Fernandes & Cordeiro, 2021; Khoo et al., 2023). Despite its industrial potential, relatively little effort has been directed toward targeted modifications of *Euglena* FA composition to enrich specific FAs (Bedard et al., 2024). A key limitation is the incomplete understanding of FA biosynthetic pathways in *Euglena*. Although some enzymes involved in this process have been characterized, such as FA desaturases FAD5 and FAD8 (Pollak et al., 2012; Wallis & Browse, 1999), many components of these pathways remain unidentified.

A complete understanding of the FA biosynthetic pathway, along with identification of the genes involved, will enable the engineering of *Euglena* strains to produce lipids with a tailored fatty acid profile, often referred to as customized lipids. Therefore, in this study, we identified the FA desaturase 12 (FAD12) enzyme, which converts oleic acid to linoleic acid in Euglena. We employed a bioinformatics approach to identify a candidate gene, and its function was investigated by overexpression in *Nicotiana benthamiana* and targeted gene knockout. As expected, *EG* FAD12 overexpression in tobacco increased linoleic acid content, while CRISPR-mediated knockout in Euglena resulted in a drastic reduction of linoleic acid. This is the first use of CRISPR in Euglena to understand FA biosynthesis and to characterize a novel FAD12 enzyme (Figure 1). The identification of the enzyme using CRISPR builds upon our knowledge of FA biochemistry in Euglena and will help open the door for customized lipid production.

**Figure 1:**
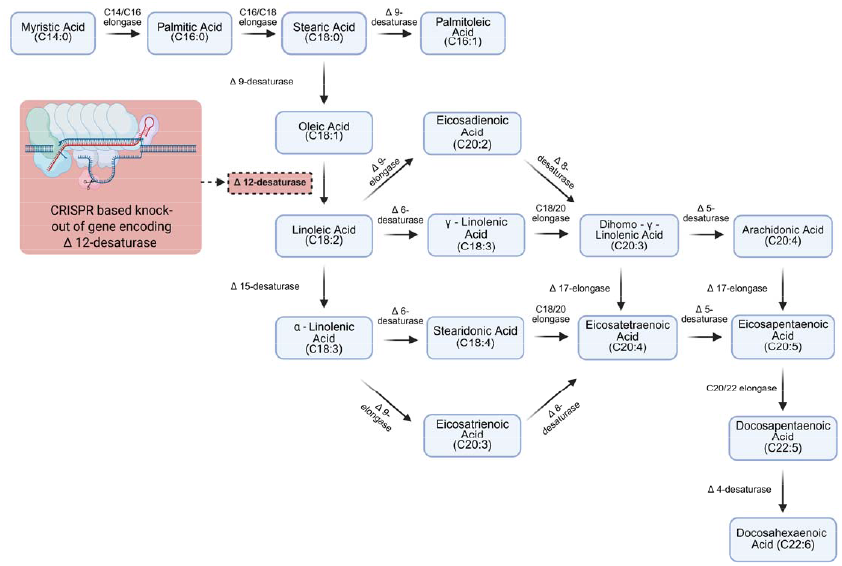
Fatty acid biosynthesis pathway. Fatty acid desaturase Δ12 is knocked out by CRISPR-Cas9 in this study.

## Methods

### 1. Organisms and growth conditions

*Euglena gracilis strain Z* (UTEX 753) was cultured in sterilized MEGM medium (sodium acetate trihydrate 1g/L, lab-lemco powder 1g/L, tryptone 2g/L, yeast extract 2g/L, calcium chloride 1g/L, and glucose 15g/L) in heterotrophic conditions as previously described (Shah et al., 2022). The cells were grown at 28ºC in a shaking incubator in the dark for a week. Then, the cells were harvested by centrifugation and used for downstream process analyses of biomass, dry mass, and lipid.

### 2. Bioinformatics Identification of the FAD12 gene in *Euglena gracilis*

A list of FAD12 proteins from 19 different microorganisms and plants was obtained from the NCBI. The HMMER software was then used to analyze the data (Eddy, 1998). First, a unique amino acid profile (HMM profile) of fatty acid desaturase protein was generated by hmmbuild, which uses the principle of the Hidden Markov Model (Supplementary Figure 1). Using the hmmsearch, the hmm profile was blasted against the translatome of publicly available RNAseq of *E. gracilis* (Mangal et al., 2022). All the hits were scanned by hmmscan for the presence of all three histidine boxes in this order (HCEGH, HXXHH, HVXHH, where X is any amino acid). Next, the transmembrane domains, which are the features of FAD12, were computationally predicted using DeepTMHMM-1.0 (Hallgren et al., 2022). Finally, we obtained one candidate protein that fits the previously described profile of FAD12. (Supplementary Figure 2).

### 3. FAD12 gene synthesis and plasmid construction for bacterial transformation

The native FAD12 transcript from Euglena was too long to synthesize by RNA isolation and cDNA synthesis, so we opted to synthesize the gene in-house using BioXP 3250 DNA printer. The synthetic gene sequence was verified by sequencing and agarose gel electrophoresis. The DNA fragment was cloned into the pEAQ-HT vector between Agel and Xmal by the restriction digestion method. The plasmid harbouring the FAD12 gene was expressed in *E*.*coli* (DH5a) by the Heat Shock method for amplification. Finally, the plasmid was transformed into *Agrobacterium tumefaciens* (GV31010) by electroporation.

### 4. Heterologous expression of *E*.*G* FAD12 in Tobacco

The transient expression of *EG* FAD12 in Tobacco (*Nicotiana Benthamiana*) was performed as previously described (Thapa et al., 2023). Briefly, the *Agrobacterium tumefaciens* (GV3101) harboring the empty plasmid (control) and the plasmid with the FAD12 coding sequence under the 35S promoter were cultured in LB broth at 28ºC with appropriate antibiotics. The bacterial cultures were centrifuged and resuspended in the infiltration medium (5mM 4–Morpholineethanesulfonic acid (MES), 50 µM acetosyringone, and 5 mM MgSO4. Then, it was injected into the ventral side of the leaves of a four-week tobacco plant. The injected portion of leaves was collected after 3 days for lipid extraction and downstream analysis.

### 5. Amino acid alignment and Phylogenetic Tree Analysis

The amino acid sequence of FAD12s from nineteen organisms was retrieved from NCBI. The amino acid alignment of twenty FAD12s and their phylogenetic analysis were conducted in the COBALT program in NCBI (Papadopoulos & Agarwala, 2007). The COBALT works by finding a cluster of pairwise constraints from database searches, sequence similarity, and user input. Then, it combines these pairwise constraints and integrates them into a progressive multiple alignment. The image of the phylogenetic analysis tree was generated from the COBALT program. The amino acid sequences from different organisms with high similarity are clustered in the same node. Those in different nodes exhibit lower similarity and indicate greater evolutionary distance.

### 6. Biomass and Lipid Quantification

To estimate the *Euglena gracilis* dry biomass weight, 5 mL of culture broth was centrifuged for 10 min at 2500 rpm. The supernatant was discarded, and the wet cells were washed twice with deionized water, dried overnight at 60°C, and weighed. Lipid was extracted using a single-step procedure as previously reported (Axelsson & Gentili, 2014). A 100 mg dried biomass sample was resuspended in a 2:1 ratio of chloroform: methanol. To achieve this, 5.26 mL of chloroform and 2.63 mL of methanol were added, and the tube was manually shaken vigorously until the biomass was dispersed entirely. Subsequently, 2.11 mL of 0.73% NaCl solution was added to produce a 2:1:0.8 (v/v/v) system, bringing the total volume to 10 mL in a 15 mL Falcon tube. The mixture was centrifuged at 4000 rpm for 10 minutes to separate the phases. The lower chloroform layer containing the extracted lipids was carefully collected in a pre-weighed vial. The extraction process was repeated with a fresh 2:1:0.8 solvent system to increase the lipid content. The combined chloroform layers were dried under vacuum to remove the solvent. Lipid content was determined gravimetrically from the weight difference.

The lipid from the tobacco plant was extracted as described (Reynolds et al., 2015). The tobacco leaves were freeze-dried and ground into a fine powder. The solvent of a chloroform: methanol mixture (2:1 v:v) was added to the powder and rotated in a shaker for 3 minutes. Then, 1:3 v:v NaCl (0.1M) was added to the mixture and rotated in a shaker for 3 minutes. Next, it was centrifuged for 5 min at 14,000 g, and a lower lipid phase was collected and dried with N2 gas.

### 7. Fatty acid analysis by Gas Chromatography Flame Ionization Detector (GC-FID)

Fatty acid methyl esters (FAMEs) of the extracted lipids were prepared using the boron trifluoride (BF_3_) methylation method as previously reported (Araujo et al., 2008) with slight modifications. Briefly, glyceryl triundecanoate dissolved in heptane was added to the lipid sample as an internal standard. Heptane was then evaporated under a gentle stream of nitrogen (N_2_), leaving the internal standard in the sample. Next, the samples were hydrolyzed with alcoholic sodium hydroxide (NaOH), followed by methylation using BF_3_-methanol solution. FAMEs were extracted with heptane, and a saturated sodium chloride (NaCl) solution was added to facilitate phase separation. The upper heptane layer containing FAMEs was transferred into a clean vial and dried over anhydrous sodium sulfate. Gas chromatography-flame ionization detection (GC-FID) was performed using an Agilent 7890B system equipped with an SP2560 column (100 m × 0.25 mm × 0.2 μm). Injection volume was one μL with a split ratio of 50:1 and a flow rate of 1.2 mL/min. Oven temperature was programmed to hold at 100°C for 5 minutes, ramp at 4°C/min to 240°C and hold for 15 minutes. Peak identification and quantification were carried out using Agilent Technologies ChemStation software.

### 8. Guide RNA (gRNA) design and CRISPR knockout of FAD12

The gRNA for FAD12 was designed using Integrated DNA Technologies software, and the CRISPR knockout experiment was conducted as described by Nomura et al., 2020). Briefly, crRNA (100 µM) and tracrRNA (100 µM) were mixed to form a complex, and then Cas9 (62 µM) was added. The CRISPR/Cas9 ribonucleoproteins (RNPs) complex was mixed with Euglena cells (1M), and electroporation was conducted. The cells were grown for 24 hours at 28ºC, and single cells were selected for further growth and analysis. The knockout of FAD12 was confirmed by DNA sequencing.

## Results

### 1. Bioinformatic Identification of FAD12 in *Euglena gracilis*

To identify FAD12, we used a bioinformatics approach based on sequence homology (Figure 2). First, we compiled a list of known FAD12 proteins from different organisms available in the National Center for Biotechnology Information (NCBI) database. Then, an amino acid profile of FAD12 (profile HMM-Supplementary Fig.1) was generated by Hidden Markov Model (Eddy, 1998). The HMM profile displays the most likely amino acid at each position of the protein, which is vital for identifying conserved and non-conserved regions. Since there were no publicly available Euglena proteome data, we took advantage of publicly available Euglena transcriptome data (Mangal et al., 2022). Using this data, we created a translated protein database of all six possible reading frames (3 Forward, 3 Reverse). The FAD12 HMM profile was searched against the translated protein data to identify candidate genes., Obtained candidates were scanned for the three conserved histidine motifs in this order (HECGH, HXXHH, and HVXHH (X=any amino acids)). Finally, we selected one candidate (532 amino acids, Supplementary Table 1) with three histidine boxes and multiple transmembrane domains, which are the peculiar characteristics of FAD enzymes. The corresponding FAD12 transcriptome sequence was aligned with our in-house Euglena genome data to identify the FAD12 genomic sequence (Supplementary Table 2).

**Figure 2:**
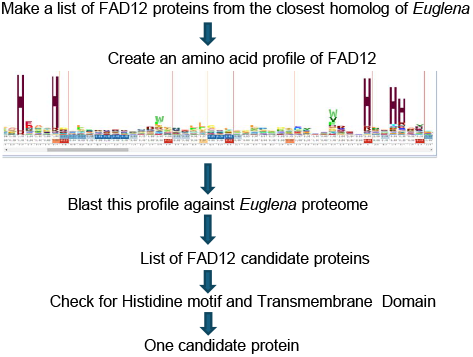
Flowchart of bioinformatic identification of FAD12 genes in *Euglena gracilis*.

### 2. Amino Acid Alignment and phylogenetic analysis of FAD12

The amino acid sequences of FAD12 from *Euglena gracilis* and other organisms were aligned using the Constraint-based Multiple Alignment Tool (COBALT), NCBI (Papadopoulos & Agarwala, 2007). The alignment focuses on conserved domains and local sequence similarity. The compact view alignment with conservative setting 2 bits is shown in Figure 3. The red letter indicates conserved regions, while blue indicates less conserved regions with no gaps. The FAD12 amino acids are vastly conserved across different organisms. The histidine boxes HECGH, HXXHH, HVXHH (X=any amino acids) are found in *EG* FAD12 along with other organisms. All the FAD12s are below 500 AA.

**Figure 3:**
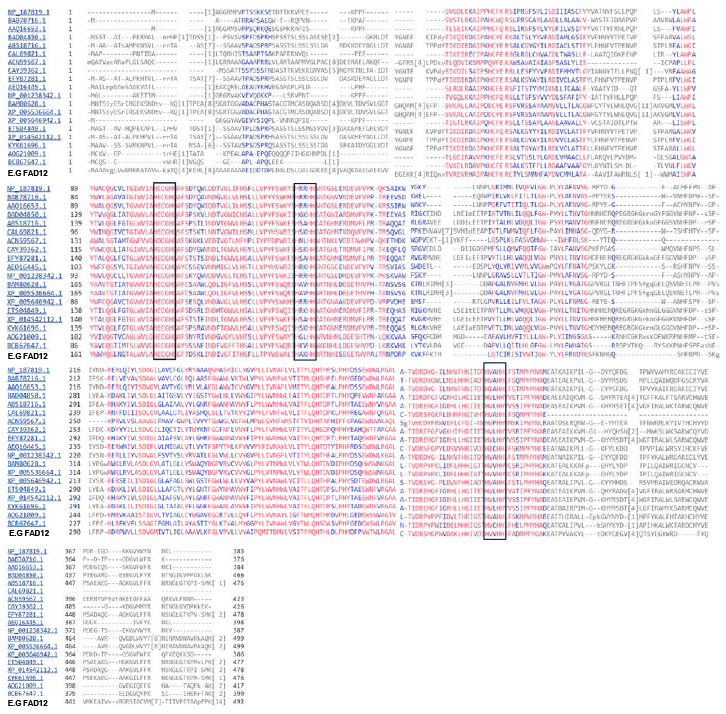
Amino acids alignment of FAD12 from different organisms. Red color indicates highly conservative regions and blue color indicates less conversed regions with no gaps. The three histidine boxes are highlighted with black boxes.

To study the evolutionary relationship between *EG* FAD12 and others, a phylogenetic tree was constructed using the amino acid sequence by the Neighbour-joining method, using the COBALT program (Figure 4). *EG* FAD12 falls into the same category as Phytophthora. It is evolutionarily nearest to Ascomycete fungi and farthest from Green Algae. FAD12 is highly conserved across species in different kingdoms, so there is high similarity (70-88%) among them.

**Figure 4:**
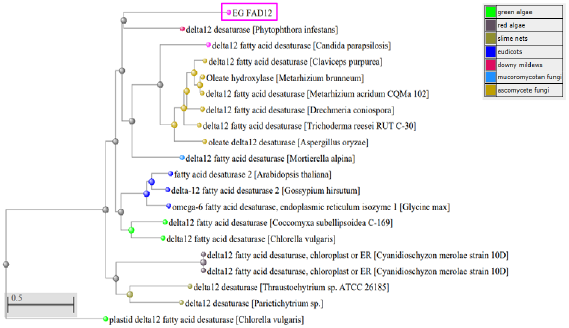
Phylogenetic analysis of FAD12 from different organisms. The phylogenetic tree was created using COBALT, max seq difference 0.85, Distance-Grishin (Protein). FAD12 from different classes of organisms are shown In different color as shown in color key. The EG FAD12 is highlighted in pink color.

### 3. Cloning of *EG* FAD12 and heterologous expression in Tobacco

Once the coding sequence of *EG* FAD12 was identified using our bioinformatics approach, we synthesized it in-house using a Codex DNA printer. Resultant DNA was cloned into the pEAQ-HT vector through the restriction digestion method and confirmed by sequencing. The size of the DNA was further confirmed using agarose gel electrophoresis (Figure 5A). The control vector and the vector harbouring FAD12 were injected into tobacco leaves, and the leaves were collected after three days for lipid extraction and analysis. Extracted lipids were methylated using the boron trifluoride (BF_3_) methylation method and were analyzed using GC-FID using the method reported by Araujo et al. (2008). There was no difference in C18:1 and C18:2 in tobacco injected with the control vector (Figure 5B). However, C18:2 levels increased in all six tobacco plant replicates injected with the FAD12 gene sequence, showing an average rise of 33%. This increase suggests an enhanced delta-12 desaturase activity. The raw data of the lipid profile are shown in Figure 5C.

**Figure 5:**
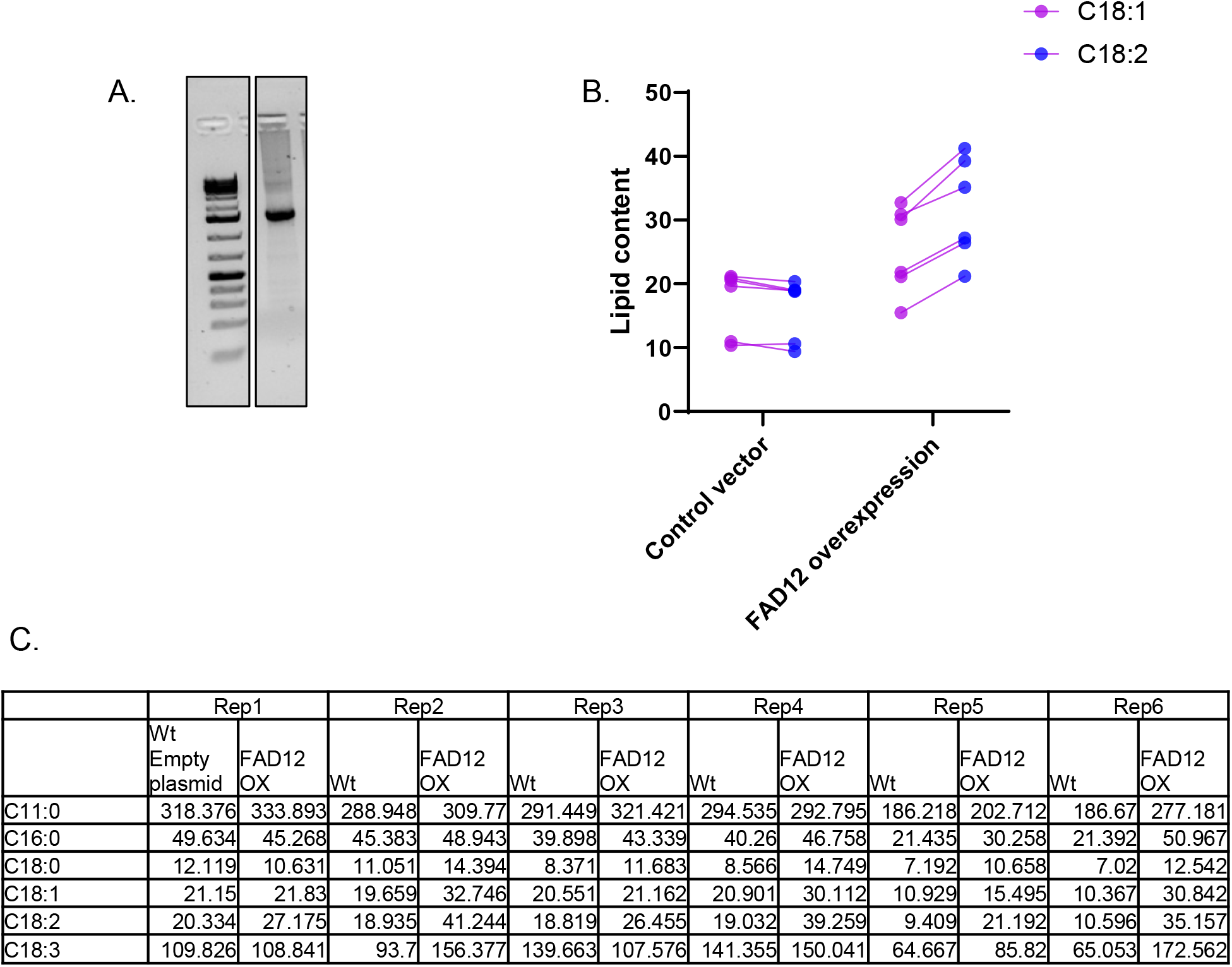
Cloning of *E*.*G* FAD12 and its heterologous expression in Tobacco. A. The coding sequence of EG FAD12 were synthesized using Codex system and ran on agarose gel to confirm the size (1476bp). B. The lipid profile (C18:1 & C18:2) of tobacco injected with control vector or FD12 overexpression vector. C. The lipid analysis data from tobacco injected with control or FAD12 OX vector.

### 4. CRISPR-Cas9 Mediated Knockout of FAD12

To further confirm the role of the identified FAD12 candidate in the fatty acid biosynthesis pathway, we used RNP-mediated CRISPR-Cas9 technology to edit this gene in Euglena. First, the FAD12 transcript was aligned with the genomic sequence to identify the gene structure, which contains 10 introns and 11 exons (Figure 6A). Then, a guide RNA was designed in the second exon to knock out the gene (gRNA and related primers are listed in Supplementary Table 3). The CRISPR-Cas9-mediated knockout was conducted as described (Nomura et al., 2020). Then, a single cell was isolated from the CRISPR knockout experiment, cultured, and sequenced to confirm the gene knockout (Figure 6B). The DNA sequencing revealed a single nucleotide deletion at position 190, which resulted in a codon shift and the appearance of multiple early stop codons (Supplementary Table 4). Under the microscope, the CRISPR-edited cell line was similar in morphology and motility to that of wild-type Euglena (Figure 6C). Lipids were extracted from this gene-edited cell line and subjected to GC-FID analysis. As predicted, compared to wild type, there was a drastic reduction in C18:2 in the gene-edited cell line (Figure 6D). The raw data of lipid analysis are available in Supplementary Table 5. Since FAD12 catalyzes the conversion of C18:1 to C18:2, knockout of this gene prevented the formation of C18:2 in Euglena.

**Figure 6:**
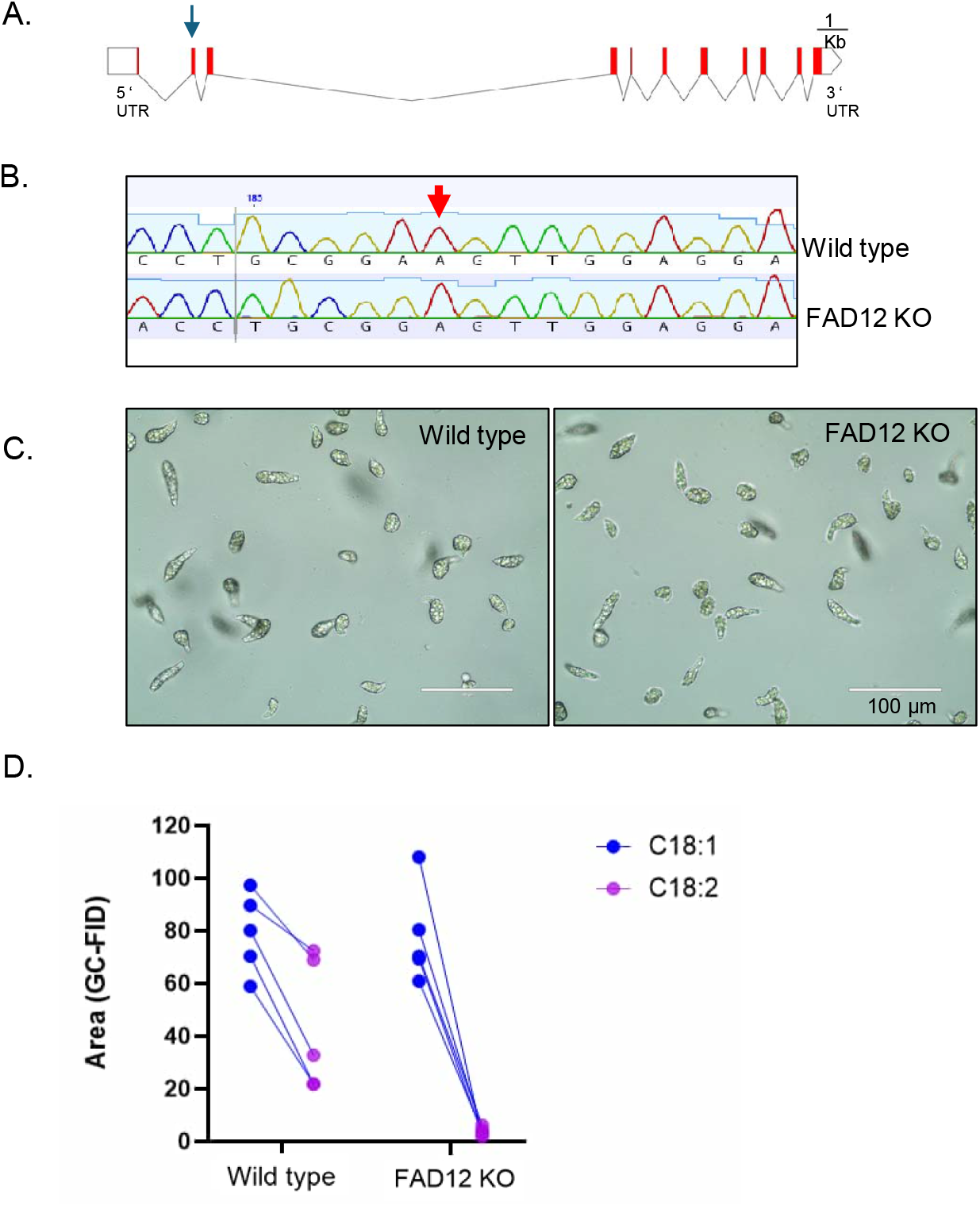
CRISPR-Cas9 Mediated Knockout of FAD12 dramatically reduced linoleic acid production. A. FAD12 gene structure and guideRNA location. B. DNA sequencing of FAD12 knockdown Euglena showing single nucleotide deletion (Adenosine) highlighted by red arrow. C. Culture of FAD12 knockdown cells selected from single cell D. Lipid profile (C18:1 & C18:2)of FAD12 knockdown cells with wild type cells

### 5. Characterization of the FAD12 kd cell line for commercial production

The FAD12 knockout cell line was further evaluated for use in commercial production. First, Euglena cells were cultured in standard growth media to compare with the wild-type strain (Shah et al.,2022). Both looked similar in color, texture, flavor, and aroma (Figure 7A). There was also no difference in total biomass and dry biomass between the two strains (Figure 7B-C). Although the lipid profile was different, there was no difference in total lipid content (Figure 7D). Finally, it was tested for glucose consumption, and both strains exhibited a similar pattern of glucose consumption over time (Figure 7E). Overall, there was no yield penalty for biomass or lipids in the FAD12 knockout strain.

**Figure 7:**
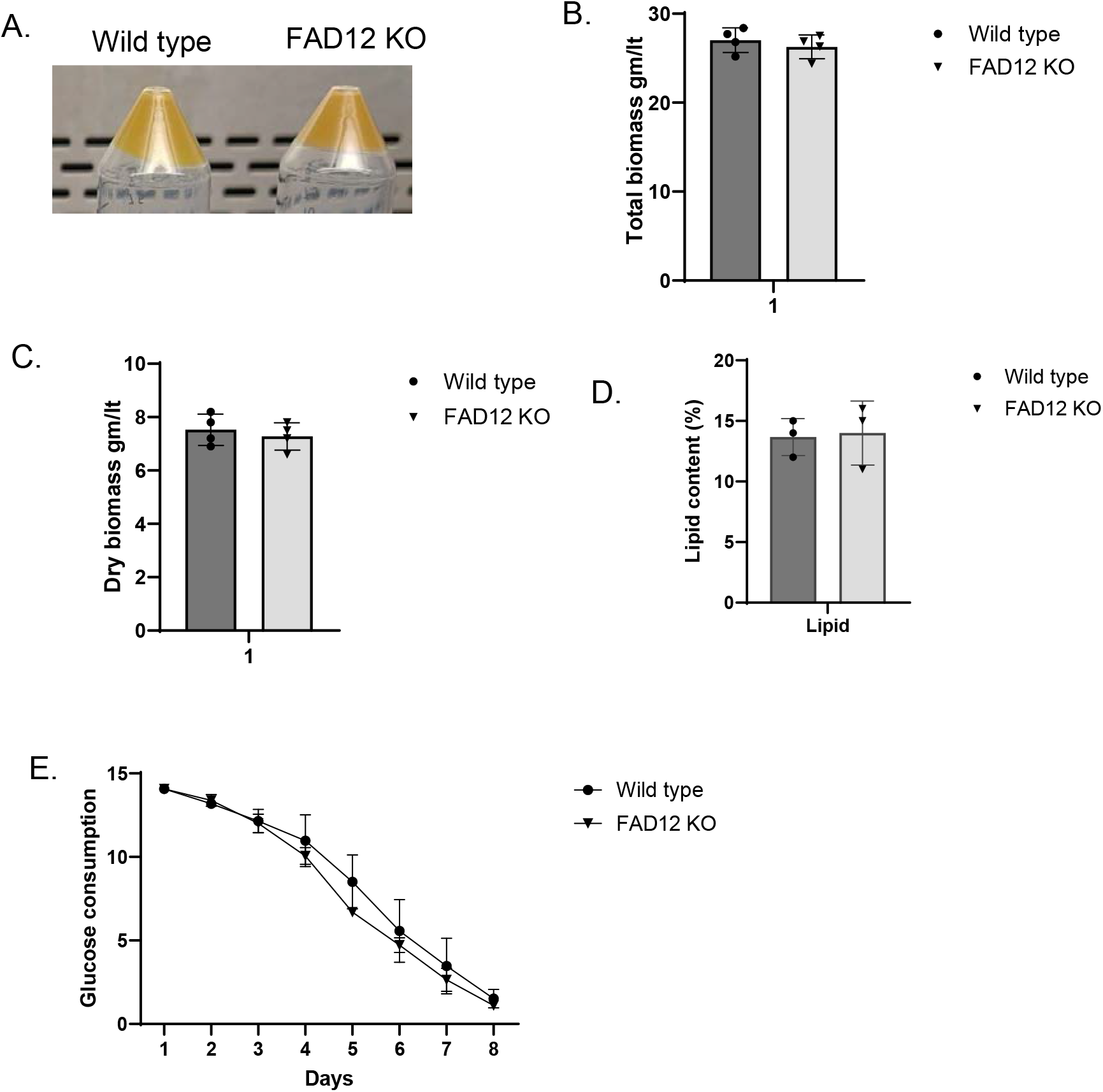
Characterization of FAD12 knockdown Euglena. A: Biomass of wildtype and FAD12Δ Euglena B: Total biomass comparison between wildtype and FAD12Δ Euglena C: Dry biomass comparison between wildtype and FAD12Δ Euglena D: Lipid content between wildtype and FAD12Δ Euglena E: Glucose consumption between wildtype and FAD12Δ Euglena

## Discussion

Euglena has emerged as a promising platform for producing high-value compounds, including tailored lipids and metabolites utilized in cosmetic, food, and nutraceutical applications (Ebenezer et al., 2022). Over the last decade, the major hindrance to studying Euglena has been the unavailability of high-quality genomic data. However, there have been some notable accomplishments recently, such as the publication of a draft genome, transcriptome, and proteome (Ebenezer et al., 2019). Also, a chromosome-level genome assembly of Euglena was published last year (Chen et al., 2024). The availability of these next-gen data is expected to expedite the understanding of several major biochemical pathways, including the fatty acid biosynthesis pathway.

To enhance lipid production in oleaginous microorganisms, including Euglena, researchers have employed both endogenous (such as pathway engineering) and exogenous strategies (including media optimization and external elicitor applications) (Matouk et al., 2025; Seemashree et al., 2022; Loh et al., 2020; Uprety & Rakshit, 2018). There has also been an attempt to co-culture Euglena with other microalgae to boost biomass and lipid production (Toyama et al., 2019). Recent efforts have particularly focused on increasing the production of specific fatty acid types in oil-producing microbes to generate customized lipids with significant industrial potential (Du et al., 2025; Duman-Özdamar et al., 2022; Wang et al., 2023). The fatty acid desaturase FAD12 represents a critical enzyme in the lipid biosynthesis pathway of oleaginous microorganisms, catalyzing the conversion of oleic acid (C18:1) to linoleic acid (C18:2). Understanding and manipulating this enzyme’s activity is of considerable interest, as it enables redirection of carbon flux to either increase or decrease specific fatty acid production (C18:1 and C18:2), ultimately facilitating customized lipid synthesis.

In this study, we employed bioinformatics, forward, and reverse genetics approaches to identify and characterize the FAD12 enzyme, a key regulator of fatty acid biosynthesis in Euglena. All characterized FAD12 proteins, including that in Euglena, possess three essential histidine domains that are crucial for desaturase activity (Supplementary Figure 2). These domains interact with iron ligands to catalyze the desaturation reaction, as previously documented (Douglas & Rousseau, 1992). The domain analysis using DeepTMHMM-1.0 (Hallgren et al., 2022) predicted that Euglena FAD12 (EGFAD12) contains five transmembrane domains (Supplementary Figure 3), a characteristic feature shared among desaturase enzymes across various organisms. The subcellular localization prediction using DeepLoc2.0 (Thumuluri et al., 2022) indicated that EGFAD12 localizes to the endoplasmic reticulum (Supplementary Table 6). This localization pattern is consistent with FAD12 enzymes from other organisms, which typically reside in the endoplasmic reticulum, the Golgi apparatus, or plastids (Dar et al., 2017; Pollak et al., 2012; Wallis & Browse, 1999). The FAD12 ER localization is expected, as the ER is one of the primary cellular sites for lipid biosynthesis.

We verified the FAD12 activity by overexpressing it in tobacco and knocking it out in Euglena using the CRISPR-Cas9 system. As hypothesized, the linoleic acid level was increased after overexpression, and the oleic acid level was increased after knockout. This complementary approach of functional gene study shows that our FAD12 candidate gene has desaturase activity. To improve its activity in the future, researchers can take the path of enzyme engineering. E.g., one can add an extra histidine motif, or replace the iron ligand with copper or other metals. Furthermore, one could optimize the enzyme expression in specific organelles for easy harvesting and downstream processing, which are critical for commercial success. Since the FAD12 KO cell line does not incur a yield penalty from altering its lipid profile, it can be further utilized for knockout or knocking in other genes and pathways.

This study presents a novel approach for customized lipid production in *Euglena gracilis* through the CRISPR-Cas9-mediated knockout. A holistic understanding of the fatty acid pathway in Euglena will help to produce a myriad of tailored lipids. Overall, these findings are expected to have significant implications for the biotechnological production of valuable lipids in microalgae for industrial applications. In particular, the production of Euglena-derived lipids enriched in oleic acid is industrially attractive due to oleic acid’s enhanced oxidative stability and its wide-ranging applications in food, cosmetics, and bio-based materials (Heggs 2022; Zambelli 2020).

## Author Contributions

Conceptualization, SCF, RJNE, BKU, and RKT; methodology, SCF, RKT, and BKU; investigation, RKT and BKU; resources, SCF and RJNE; data curation, RKT and SCF; writing, original draft preparation, RKT; writing, review, and editing, RKT, BKU, RJNE, and SCF; supervision, RJNE and SCF; project administration, RJNE and SCF; funding acquisition, RJNE and SCF. All authors have read and agreed to the published version of the manuscript.

## Acknowledgement

This work was funded by Noblegen and MITACS. Raj Kumar Thapa and Bijaya Kumar Uprety were recipients of the MITACS Accelerate Industrial Post-doctoral award. We would like to thank the Cell & Systems Biology team from Noblegen and the Emery lab for experimental assistance and helpful discussions.

---

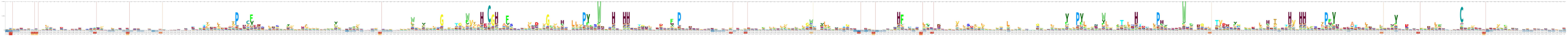

**Supplementary Figure 2:**
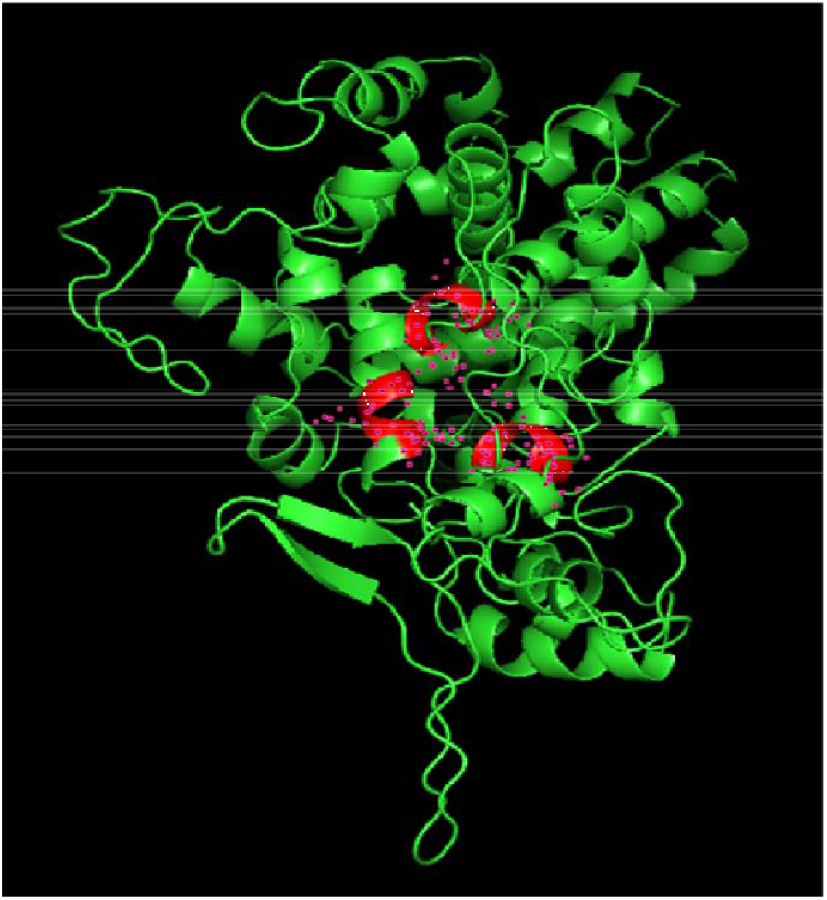
Computational prediction of the 3D structure of EG FAD12. The image was generated using PyMOL. The red color indicates the histidine box in FAD12, which comes together to interact with iron.

**Supplementary figure 3:**
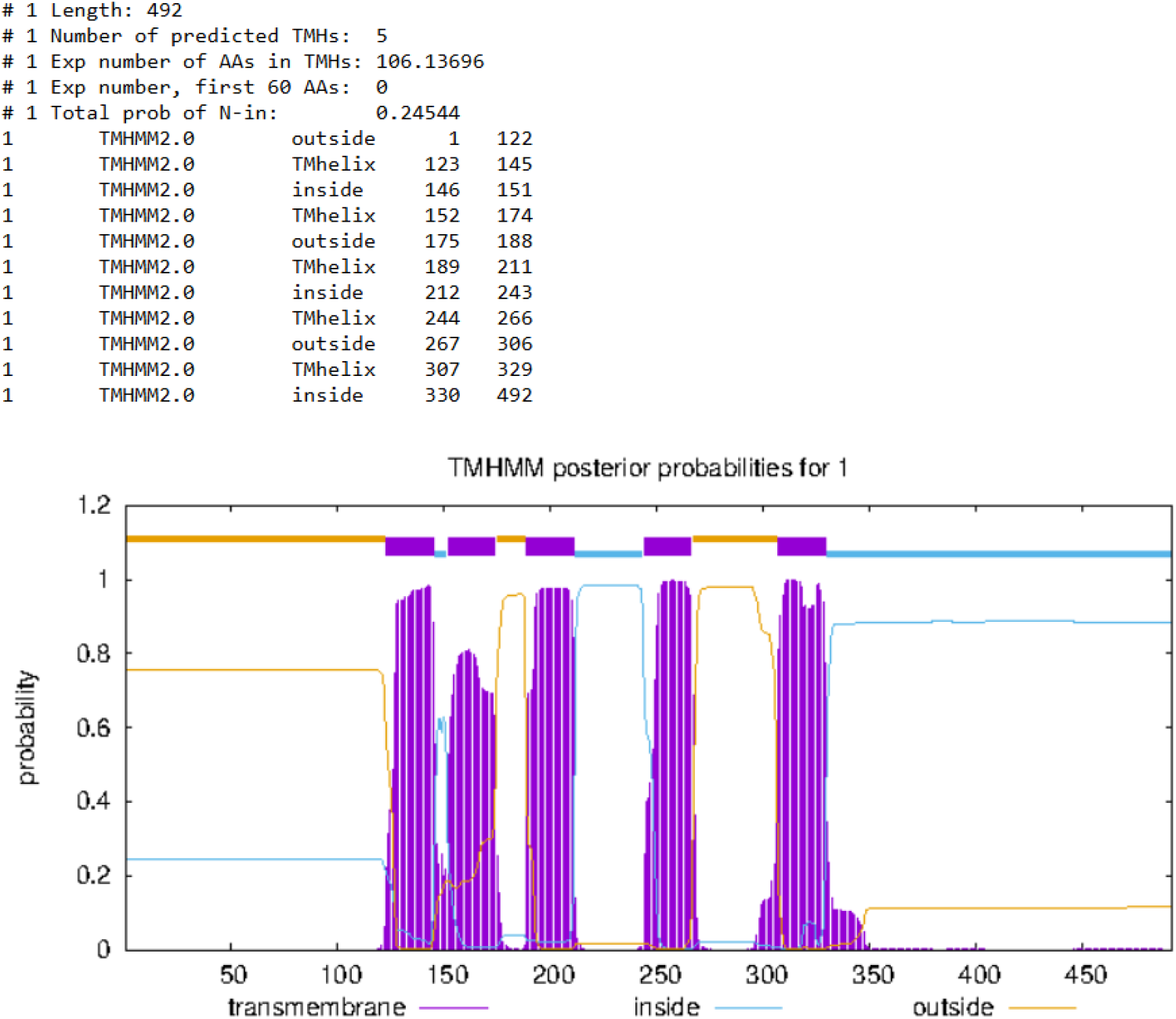
Prediction of transmembrane domains (TMD) in EG FAD12 by TMHMM - 2.0. It is predicted to have five TMDs in different positions as shown in the figure.

## References

Araujo, P., Nguyen, T.-T., Frøyland, L., Wang, J., & Kang, J. X. (2008). Evaluation of a rapid method for the quantitative analysis of fatty acids in various matrices. Journal of Chromatography A, 1212(1), 106–113. 10.1016/j.chroma.2008.10.006

Axelsson, M., & Gentili, F. (2014). A Single-Step Method for Rapid Extraction of Total Lipids from Green Microalgae. PLOS ONE, 9(2), e89643. 10.1371/journal.pone.0089643

Bedard, S., Roxborough, E., O’Neill, E., & Mangal, V. (2024). The biomolecules of Euglena gracilis: Harnessing biology for natural solutions to future problems. Protist, 175(4), 126044. 10.1016/j.protis.2024.126044

Chen, Z., Chen, Y., Zhang, H., Qin, H., He, J., Zheng, Z., Zhao, L., Lei, A., & Wang, J. (2022). Evaluation of Euglena gracilis 815 as a New Candidate for Biodiesel Production [Original Research]. Frontiers in Bioengineering and Biotechnology, Volume 10 - 2022.

Chen, Z., Dong, Y., Duan, S., He, J., Qin, H., Bian, C., Chen, Z., Liu, C., Zheng, C., Du, M., Yao, R., Li, C., Jiang, P., Wang, Y., Li, S., Xie, N., Xu, Y., Shi, Q., Hu, Z., … Wang, J. (2024). A chromosome-level genome assembly for the paramylon-producing microalga Euglena gracilis. Scientific Data, 11(1), 780. 10.1038/s41597-024-03404-y

Dai, J., He, J., Chen, Z., Qin, H., Du, M., Lei, A., Zhao, L., & Wang, J. (2022). Euglena gracilis Promotes Lactobacillus Growth and Antioxidants Accumulation as a Potential Next-Generation Prebiotic. Front Nutr, 9, 864565. 10.3389/fnut.2022.864565

Dar, A. A., Choudhury, A. R., Kancharla, P. K., & Arumugam, N. (2017). The FAD2 Gene in Plants: Occurrence, Regulation, and Role [Review]. Frontiers in Plant Science, Volume 8 - 2017.

Douglas, R. G., & Rousseau, D. L. (1992). Hydrogen bonding of iron-coordinated histidine in heme proteins. Journal of Structural Biology, 109(1), 13–17. 10.1016/1047-8477(92)90062-F

Du, F., Zhang, F., Hang, Y., Jing, H., Zheng, Y., Ma, W., Sun, X., & Huang, H. (2025). Advances in production of customized functional unsaturated fatty acids in Yarrowia lipolytica. Agricultural Products Processing and Storage, 1(1), 14. 10.1007/s44462-025-00016-6

Duman-Özdamar, Z. E., Martins dos Santos, V. A. P., Hugenholtz, J., & Suarez-Diez, M. (2022). Tailoring and optimizing fatty acid production by oleaginous yeasts through the systematic exploration of their physiological fitness. Microbial Cell Factories, 21(1), 228. 10.1186/s12934-022-01956-5

Ebenezer, T. E., Zoltner, M., Burrell, A., Nenarokova, A., Novák Vanclová, A. M. G., Prasad, B., Soukal, P., Santana-Molina, C., O’Neill, E., Nankissoor, N. N., Vadakedath, N., Daiker, V., Obado, S., Silva-Pereira, S., Jackson, A. P., Devos, D. P., Lukeš, J., Lebert, M., Vaughan, S., … Field, M. C. (2019). Transcriptome, proteome and draft genome of Euglena gracilis. BMC Biology, 17(1), 11. 10.1186/s12915-019-0626-8

Ebenezer, T. E., Low, R. S., O’Neill, E. C., Huang, I., DeSimone, A., Farrow, S. C., Field, R. A., Ginger, M. L., Guerrero, S. A., Hammond, M., Hampl, V., Horst, G., Ishikawa, T., Karnkowska, A., Linton, E. W., Myler, P., Nakazawa, M., Cardol, P., Sánchez-Thomas, R., … Field, M. C. (2022). Euglena International Network (EIN): Driving euglenoid biotechnology for the benefit of a challenged world. Biology Open, 11(11), bio059561. 10.1242/bio.059561

Eddy, S. R. (1998). Profile hidden Markov models. Bioinformatics, 14(9), 755–763. 10.1093/bioinformatics/14.9.755

Farjallah, A., Fillion, M., & Guéguen, C. (2024). Metabolic responses of Euglena gracilis under photoheterotrophic and heterotrophic conditions. Protist, 175(3), 126035. 10.1016/j.protis.2024.126035

Fernandes, T., & Cordeiro, N. (2021). Microalgae as Sustainable Biofactories to Produce High-Value Lipids: Biodiversity, Exploitation, and Biotechnological Applications. Mar Drugs, 19(10). 10.3390/md19100573

Hallgren, J., Tsirigos, K. D., Pedersen, M. D., Almagro Armenteros, J. J., Marcatili, P., Nielsen, H., Krogh, A., & Winther, O. (2022). DeepTMHMM predicts alpha and beta transmembrane proteins using deep neural networks. bioRxiv, 2022.2004.2008.487609. 10.1101/2022.04.08.487609

Khoo, K. S., Ahmad, I., Chew, K. W., Iwamoto, K., Bhatnagar, A., & Show, P. L. (2023). Enhanced microalgal lipid production for biofuel using different strategies including genetic modification of microalgae: A review. Progress in Energy and Combustion Science, 96, 101071. 10.1016/j.pecs.2023.101071

Kim, S., Im, H., Yu, J., Kim, K., Kim, M., & Lee, T. (2023). Biofuel production from Euglena: Current status and techno-economic perspectives. Bioresource Technology, 371, 128582. 10.1016/j.biortech.2023.128582

Ko, Y., Baek, H., Hwang, J.-H., Kim, Y., Lim, K.-M., Kim, J., & Kim, J. W. (2023). Nonanimal Euglena gracilis-Derived Extracellular Vesicles Enhance Skin-Regenerative Wound Healing. Advanced Materials Interfaces, 10(4), 2202255. 10.1002/admi.202202255

Kuhne, A. M., Morrison, E. N., Sultana, T., Kisiala, A. B., Horlock-Roberts, K., Noble, A., & Emery, R. J. N. (2023). Cultivation of heterotrophic Euglena gracilis: The effects of recycled media on culture growth and associations with growth regulating phytohormone profiles. Journal of Applied Phycology, 35(5), 2161–2175. 10.1007/s10811-023-03062-4

Lewis, A., & Guéguen, C. (2020). How growth conditions of Euglena gracilis cells influence cellular composition as evidenced by Fourier transform infrared spectroscopy and direct infusion high-resolution mass spectrometry. Journal of Applied Phycology, 32(1), 153–163. 10.1007/s10811-019-01929-z

Mahapatra, D. M., Chanakya, H. N., & Ramachandra, T. V. (2013). Euglena sp. as a suitable source of lipids for potential use as biofuel and sustainable wastewater treatment. Journal of Applied Phycology, 25(3), 855–865. 10.1007/s10811-013-9979-5

Mangal, V., Donaldson, M. E., Lewis, A., Saville, B. J., & Guéguen, C. (2022). Identifying Euglena Gracilis Metabolic and Transcriptomic Adaptations in Response to Mercury Stress [Original Research]. Frontiers in Environmental Science, Volume 10 - 2022.

Matouk, A. M., Abu-Elreesh, G. M., Abdel-Rahman, M. A., Desouky, S. E., & Hashem, A. H. (2025). Response surface methodology and repeated-batch fermentation strategies for enhancing lipid production from marine oleaginous Candida parapsilosis Y19 using orange peel waste. Microbial Cell Factories, 24(1), 16. 10.1186/s12934-024-02635-3

Nguyen, N. H., Nguyen, Q. T., Dang, D. H., & Emery, R. J. N. (2023). Phytohormones enhance heavy metal responses in Euglena gracilis: Evidence from uptake of Ni, Pb and Cd and linkages to hormonomic and metabolomic dynamics. Environmental Pollution, 320, 121094. 10.1016/j.envpol.2023.121094

Nomura, T., Yoshikawa, M., Suzuki, K., & Mochida, K. (2020). Highly Efficient CRISPR-Associated Protein 9 Ribonucleoprotein-Based Genome Editing in Euglena gracilis. STAR Protocols, 1(1), 100023. 10.1016/j.xpro.2020.100023

Ochsenreither, K., Glück, C., Stressler, T., Fischer, L., & Syldatk, C. (2016). Production Strategies and Applications of Microbial Single Cell Oils [Review]. Frontiers in Microbiology, Volume 7 - 2016.

Papadopoulos, J. S., & Agarwala, R. (2007). COBALT: constraint-based alignment tool for multiple protein sequences. Bioinformatics, 23(9), 1073–1079. 10.1093/bioinformatics/btm076

Park, S.-y., Kim, K. J., Jo, S. M., Jeon, J.-Y., Kim, B.-R., Hwang, J. E., & Kim, J. Y. (2023). Euglena gracilis (Euglena) powder supplementation enhanced immune function through natural killer cell activity in apparently healthy participants: A randomized, double-blind, placebo-controlled trial. Nutrition Research, 119, 90–97. 10.1016/j.nutres.2023.09.004

Pollak, D. W., Bostick, M. W., Yoon, H., Wang, J., Hollerbach, D. H., He, H., Damude, H. G., Zhang, H., Yadav, N. S., Hong, S.-P., Sharpe, P., Xue, Z., & Zhu, Q. (2012). Isolation of a Δ5 Desaturase Gene from Euglena gracilis and Functional Dissection of Its HPGG and HDASH Motifs. Lipids, 47(9), 913–926. 10.1007/s11745-012-3690-1

Reynolds, K. B., Taylor, M. C., Zhou, X.-R., Vanhercke, T., Wood, C. C., Blanchard, C. L., Singh, S. P., & Petrie, J. R. (2015). Metabolic engineering of medium-chain fatty acid biosynthesis in Nicotiana benthamiana plant leaf lipids [Original Research]. Frontiers in Plant Science, Volume 6 - 2015.

Rodríguez-Zavala, J. S., Ortiz-Cruz, M. A., Mendoza-Hernández, G., & Moreno-Sánchez, R. (2010). Increased synthesis of α-tocopherol, paramylon and tyrosine by Euglena gracilis under conditions of high biomass production. Journal of Applied Microbiology, 109(6), 2160–2172. 10.1111/j.1365-2672.2010.04848.x

Shah, M. R., Morrison, E. N., Noble, A. J., & Farrow, S. C. (2022). A simple and effective cryopreservation protocol for the industrially important and model organism, Euglena gracilis. STAR Protocols, 3(1), 101043. 10.1016/j.xpro.2021.101043

Shibata, S., Arimura, S. I., Ishikawa, T., & Awai, K. (2018). Alterations of Membrane Lipid Content Correlated With Chloroplast and Mitochondria Development in Euglena gracilis. Front Plant Sci, 9, 370. 10.3389/fpls.2018.00370

Thapa, R. K., Tian, G., Lu, Q. S. M., Yu, Y., Shu, J., Chen, C., Song, J., Xie, X., Shan, B., Nguyen, V., Li, C., Bian, S., Liu, J., Kohalmi, S. E., & Cui, Y. (2023). NUCLEOPORIN1 mediates proteasome-based degradation of ABI5 to regulate Arabidopsis seed germination. bioRxiv, 2023.2008.2010.552853. 10.1101/2023.08.10.552853

Thumuluri, V., Almagro Armenteros, J. J., Johansen Alexander R., Nielsen, H., & Winther, O. (2022). DeepLoc 2.0: multi-label subcellular localization prediction using protein language models. Nucleic Acids Research, 50(W1), W228–W234. 10.1093/nar/gkac278

Toyama, T., Hanaoka, T., Yamada, K., Suzuki, K., Tanaka, Y., Morikawa, M., & Mori, K. (2019). Enhanced production of biomass and lipids by Euglena gracilis via co-culturing with a microalga growth-promoting bacterium, Emticicia sp. EG3. Biotechnology for Biofuels, 12(1), 205. 10.1186/s13068-019-1544-2

Uprety, B. K., Morrison, E. N., Emery, R. J. N., & Farrow, S. C. (2022). Customizing lipids from oleaginous microbes: leveraging exogenous and endogenous approaches. Trends in Biotechnology, 40(4), 482–508. 10.1016/j.tibtech.2021.09.004

Uprety, B. K., & Rakshit, S. K. (2018). Use of Essential Oils From Various Plants to Change the Fatty Acids Profiles of Lipids Obtained From Oleaginous Yeasts. Journal of the American Oil Chemists’ Society, 95(2), 135–148. 10.1002/aocs.12006

Wallis, J. G., & Browse, J. (1999). The Δ8-Desaturase ofEuglena gracilis:An Alternate Pathway for Synthesis of 20-Carbon Polyunsaturated Fatty Acids. Archives of Biochemistry and Biophysics, 365(2), 307–316. 10.1006/abbi.1999.1167

Wang, J., Yu, X., Wang, K., Lin, L., Liu, H.-H., Ledesma-Amaro, R., & Ji, X.-J. (2023). Reprogramming the fatty acid metabolism of Yarrowia lipolytica to produce the customized omega-6 polyunsaturated fatty acids. Bioresource Technology, 383, 129231. 10.1016/j.biortech.2023.129231

Wang, Y., Seppänen-Laakso, T., Rischer, H., & Wiebe, M. G. (2018). Euglena gracilis growth and cell composition under different temperature, light and trophic conditions. PLOS ONE, 13(4), e0195329. 10.1371/journal.pone.0195329

