## Supplemental Table 1 for "Identification and characterization of the delta-12 fatty acid desaturase from *Euglena gracilis*"

EG FAD12 protein

MAAAVGLVWNRKRATAVWKATQERRVKEHLRKLEESKQAAAKAAAAEEDDTDPLDALDSLDALGDKKETSSLRARKVAQKAAEGAEKVVEGEKKSVLERIQNVTVREVRHAIPAHCFKRCTYLSMMHFAYDVCMMLATAAAVIFAWRNLSGWWMAIIWPAYWWYQGLNGTALWVLAHECGHGGFTDSKLINDIVGFITHSFLLTPYWSWALTHAKHHRRTNHISEGETWVPAITRNPDRPKVKFFKTHLGTCIRISIVWTIGWYLYLFRNDTGSFKNHGQSHFNPNSKGLFRPADRPWVLLSDVGMIVTLIGLAAAVLKFGLVPVLLVYAIPQMITDMYLVSITFMQHTHLDLPHMDFPVWTWLNGALCTVDRSMGPWVDAKLHHIVDTHVVHHIFADMPFYGAKEATPYVAKYLTDKYGEGVYNSKQTSYLGYWRDFYQVMKEAIVVVRGRSIACVHPHPCPCLTIIVPETAAPPPKAAFLHIFGWSHCSR
