## Supplemental Table 2 for "Identification and characterization of the delta-12 fatty acid desaturase from *Euglena gracilis*"

| Gene | scaffold | start | stop | strand |
| --- | --- | --- | --- | --- |
| D12Can1 | tig00001963_1_f_1 | 309463 | 343976 | ++ |

Coding sequence start :ATGGCAGCCGCTG

GTTATTATATTATTATCAATTTTATTAATATATATTTTCTATATTTATATAAATTTTATTATTATCAATTTTTAATGTAATATTTTATAATACAAACAGCACACGTTTTTATACAATTTTAGCAATAATATATACACACACACACACACATATATATATATAATATAGAAGTTGACATATAATGACGTCTAGAAATTGAAATTAATAAATAATTAAGTCAAAATAGTATTGATGTAATATGTAATTGTTAATGATAATTATCATGACCACTTCAAATCATATTATTGATAAAATTGAGTATGATAAAATATATGAATATGATAAAATTGAATATGATAAAAATAAAATATGATAGAATGTAAAATATGTTATTGGTAATGGGTACAGGATTATGTTCACATATTTCACTTTTTATTGTTTTTGGGAGTAATTTTTATTATTTTATTATTTTAGTATTTTATTATTTTATTTTGTCTAAAACAATGCCAAAATCAAATCGAATTTCACCAGGCAACAGCATCATCTTTCAGAAAAAAATGACTTCTTATTGTGAAACCCCGCAAAATGGTATATTTTTAAAAAAATACAAAAAACCGGGAGCCCTTCTGGTAGTGCAAATAAAACTTTTTAATATTTGTTCAACGTTTTTTTTCCTGAAAAATTTTCACTGTCGTAAATTCTTGCTGTTCAGAATTCCTCTATGGGAATGAATTAAGATTTGTTGAGTTTCCCATGGTTTTGATCCAGTGTTCATGAATTTTGCATGCTTTTTTGGATGTACATTTGTTACAACTGAATAAGAGAAGGTTTGCACCTACGTTATGATACAGTTTTCGGGACAAAAATAACGAAAAAATATGATTTTCAGAACCCCCACCTATACCTTAAAATGGCCTAAAATGGCCTTAAATGGCCTAAAAAGGCCTAAAACGGCCTAAAATGAGGAAATTTTTGGGCTTGACTAAGGTGCCTGGTCCAGACCGACCCTGGGGGGGGGGTACCCAGGACTCCCCCCAAAAGGGTCAATTTTTTCCTCCGGCGCTTTGCGCCGAAAATTTTTTGCATTTTGGCAATTTTTTTTTGCCCAAAAAGTGCCCAAAATGGCACTTTTGGTCGTTTTGGGGGGTCAAAAAATTTCGGTCACCTGGACCGCGCCACTGTTTAGGTGAGGGGTGTCCCGCAGGGGGGGGGGGTCCGGACCACCCCCCCCCCCCCACCTGTCCTTATATCAAGCCCTGCCAGGCCCCCATTAGGCTATTTTTTTGGCTTTTTTTCCCCGCTCGAAGCACCAAATTAATTGACCCTAAAATTGAATTTGAAAATGGAATTAGTTTTTATAAAATGGGTCTTTCCCCAGTCTGGGTGGCCAATTTCACAATGGTTGCCGGTGGACACACCCGAGCCCAGCACCCCCCCCACCCCAGGCCCAAGACAGGGGGCTCCGAACCCGCAATAGGTAATAGAAAATAGGAGGAGGTTGTCTGCTATTCAAATTCCTCCCCCTTGGGAACCGTTTGCTCTCTTCTTTTCCAGTTCAACCCAACCAAGCTGGTTCTGGACATGGCAGCCGCTGTGGGATTGGTTTGGAACCGCAAGCGCGCCACAGGTGTTTCCTGCTGAGACGGATGCAGCTGGTGTTTACGCTTTGAAGACCATTCACCCGCACCAGCAAACCAAATTATGAATTTTTATTGTCAGCTGTCCCGGGCAATTGAGGCATTGTCGCTTTTCAGGACCGAAATGTTGGCGACGGTTCTAGGCCTTTGATGATTTTTTTTCTCAGAAACTGGGAAGAGTAAGTCAATTTAATTGGGGCAAAATCAAAAACGTCTGGAATTCCCAAACCAATGTTTCTGGGACACTGCAAAATCAGGAGTTTTCTCAAGCTGAAGGGAAGCTGAAGGGGGGGGTCCAGCAAGCCAGTTCTTTTTCCTCAGCCACCCGGGGGGGGGGGCGCTAAAATAATTTTCAAACCAGCCCTTCAAAACCCGGACTGAAAATTTCGACACAAAGCGCCGAATATTTTTGGGAAAATGGTTTAAAATGTAATTTTGGGTGGTTTTTGGGTCCAACTGCAAATCTTAGCAGAAAAATGCCAGCCCAACTGGATTTGGGGATTTTTTTTGCTAGTTTTTGGGTGTTTGCTGGGTCCAGCTGCCATCGCCTTTTTCCAATGTTCAAAATTGAATTCAAGCTTGGCTTTCCGGCCCAAAACAGAGCAGTTCCGAACCAGCTTTCTTTTAAGGCTGCACCGCCAATAAGGAAAGAAGGCAGGGTTTTTCCTTAAAGGTTGAGCCCGTGGAGGGTGGATGGGTCTGCTGGAACCCCCCCCCCCCCCGAGCCTTGTATGGGTCCCGCAGTTAATGCAGGTTTGGCCATTTTTTGAATTTTTTTTCTGGTGCAAAGCACCAAAAAAATTTGGCCTCAGCATTACCCCAAAAATGGATCCCTGTGGGAGGGGGAAGGGGTCCCCTGGGTGGGTTGGGGACCTCCCTCCCACGTCTGAAAAAAAGAATCCCTGCAGGAAGTCCAGCCGGGACCCAAACTGTCTCTTGCGGTAGTGGTTCGTGATCAGCTGTTTTTTTGATAATTTGGTGATCCAGGGTTTTGGTCGGTGGGTTCACCCCAGGGTTCGATATAAGGAGGGCGTACATGGTTTAGGTGGGGGGGGTGTCCCCCCCACCCCCCCCCCCCCTCCTGGCCCGCAGGGGGGGGTCCGGACCACCCCCCCCCCCACCTGCAAGGGGGGTCCGGACCGACCCTCGGGGGCAAAAATTGACCCCCCCAAAAAAGGGTCAATTTTTTCCTCCGGCGCTTTGCGCCGAAATTTTTTTGCATTTTTGGCACATTTTTGCCCAAAAAGTGCCCAAAATGGCACTTTTGGTCGTTTTGGGGGGTCAAAAATTTTCGGCCACCCTGACCGCGCACCTGTTTAGGTGGGGGGGGTGTCCCCCCCCACCCCCCCCCTCCTGGCCCGCAGGGGGGGGTCCGGACCACCCCCCCCCACCTGTCCTTATATCGAACCCTGGTTCACCCCGAACGCAAGGGGTTCCTGTGGGGGGGGGGTGCGCCTGACCCCCGGGTCCGAAAAAACTTTGTCAGCAGAAGGACCCTGGGTGATATGTGGTTTAGGCTTGCTCAGAACGGATTCACCTGCATTTGAGTGTTCAACATTTCTAGGGACTGCGCAAAGAGATTTTGCCTCATTTCAAATTGTTGTTGAAAATTATCTCCATTGTGATTTATTCACATGTCCGTTTAGCAAGCAGCATTTTTTTTGATACTGCCGGGTCAGAGCGAGGGCAATGGGACATGGCAACCCTTCTCCATCAACCCCCCTCTTGCAGCGGTGTGGCAGGCGACGCAGGAACGGCGCGTCAAGGAGCACCTGCGGAAGTTGGAGGAGAGCAAGCAGGCGGCGGCCAAGGCCGCGGCGGACAGGGCCGTATCGGAATCCCGGAAACCCTGTGGTGGGTGGGTGCCCCCCCCCTGTCCCCATCCGGTGGGGGCAGGGCAGAACGGGAATCAGCCAGTCAGCCAGGGTGGTGGGGGGGACCCCCCCCCACCGCTCCCGTAATTCCGGGAGCACCAATTTTCGACCCCAAAATTTTTTTCCCGGCGCCAGGGAAAATTTGGTTAAAAATCTGTGAAAATGCTGTGGTGGGTGGGAGGGTCAACCACCCCCACCTCCCAGATGGGGGGGTCGACCACCCCACCAGGCTTTCCCGTTTTGCCCTGGGTGGGCGCTCCCCATATTTCTGGGCGTGAATTTTTGTGTCAAAAGTTTCTGTCGGCGCCGAAGGTTTTTTCCTTCCTGGGGGGTGACCATCCCCCCAGGCTTCGTTGTGCAGCCCTGGCAGAGGCGGAGGAGGACGACACGGACCCCCTGGACGCCCTGGACAGCCTGGACCCCCTGGGGGAGAAGAAGGAAACCTCTTCCCTCCGGGCCCGCAAGGTTGCGCACAAGGCCGCGGAGGGGGCGGAGAAGGTGGTGGAAGGGGAGAAGAAAAGCGTTCTGGTGTGTCCACCGATCGTCTGCTGACAGCCAACCTGGCAACATCTCTTCCATTGAGACCCAAAAGTGTGTGGTTTATCACCTTCACCTGGGATGTTGGTTTTTCTTGCAGCTGCAATTTTGGAGTACTCCAGGGTGTATTCAAGGTATTCCCGTGACGGGTAGGGAAGGGTCTGGTTCGGAATGTCCCCTCCCCCCGGAGGCCCCACCCTTGCACCAGGAGCCAGAATCGCACCCAAAAAATATTCTGGCCTTTTTTTTTGCATTTTGCATTTTTGCATTTTTTTTTTTAGTTTGGGAAAGGTCCCAAATACTTGACACGCACTCGTAATTTTAATTCGAGGGATTTTGACGGTTTTTTTTTAGAAAGAATCCTCAAATGAAGGCCTCACTGGGGGAGGGAGTGGGCATCCAAAGGGTTATCCCGTTTTGGCCGAAGACATCAGCCCATTTTTGAAATATTTTTTTGCACACATGCTGCGGTAAGTTGCAACTGTACATTCACTGAACCTCTGTGCATCCCCTCGCCTTTTAGAGTGATTTGTGTTGTGGGTGTTATGGGCAGGCAGGGGGTTTGGCATTTCGACCTCCCCCCCCCCGCAGGGGAGGGATCGTGTTTTGGGGTGGGCAGTGGTATAGCACACCGTCGCTGCTGAAACAGGCCGGGTCCCTAAAGTGGCCACTTCAAAAGGGAGAATTGGCTCAGTCGGAAGGCACCAGTGCCCAAGGCCGAGCCCCGCACCCGGGTAGGCACCCCATAGCAGAAGGCACCCCCCCGTCCTCCCCGGGGGCTGGGGGGTGGGGGGCTGGCGTGACCAAGTCGGAGGGTTGTGTGGGTGGGTTATCCAATGAAAGTGCATGGGCCCCAATGCCCATCTTTGCAACCATCCACCGTCGTCTTTGCCAGTCCCGTCTGACCCGCATGGCTGGGCTGGTGGGTAGAGGTGGAGAGATGTATGCGCAAATGGCATTTTGTTGGCAGGTCTCCTCTCCTCTGGCACCTGCATCGGGCCCCAGGGTTTGGTTGCTTGCCCGTCAACCGTTGGGTTTGAAGGGGAGGGGTATTCTCCGAACGGGAACCAGCCCGCTTGGTGGCGTTTCGCATGATGACCCCCTTCTCAGCGGGTTCCACTGATTAGTGATCGAGAGTGACAACCGAGCGACTGAGAGCGATGGCTGGTGATGTGTGCTCGTGCTCCATGGGCAGCGTTGGGTGGGAGGCCTGGATGGTGTCACATGATGTACTGATTGTTCAATTCCTTTACCTGCACCGTGATGAACATTCCCAAGCTGATGGGTGCTGGGTGGTTGGAAGCACTGCGGAAACACACCACTCGTGTGGCAGAGGCACTTGGTAACCTGGGATGCGAGGCCTTGAGTGATGATTCCTCTCAGCGGGTTTCTGAGACGATGCTGAGGCTCCATAATTCTGGCTGATAAGGGCGGGAAGCAACTGGTGATTGGGGTGTGTCCAGCAGCAGGGGTTCGGAGGGTAGTGGGTGCGGTAAGTTGCAACTGTACATTCACTGAACCTCTGTGCATCCCCTCGGTGATTTGTGTTGTGGGTGTTATGGGCAGGCAGGGGGTTTGGCATTTCGACCCCCCCCCCCCCCCGCAGGGGAGGGATCGTGTTTTGGGGTGGGCAGTGGTATAGCACACCGTCGCTGCTGAAACAGGCCGGGTCCCTAAAGTGGCCACTTCCAAAAGGGAGAATTGGCTCAGTCGGAAGGCACCAGTGCCCAAGGCCGAGCCTCGCACCCGGGCAGGACCCCATACATAGCAGAAGGCACCCCCGTCCTCCCCGGGGGCTGGGGGGGGGCTGGCGTGACCAAGTCGGAGGGTTGTGTGGGTGGGTTATGAAAGTGCATGGGCCCCAATGCCCATCTTTGCAACCATCCACCGTTGTCTTTGCCAGTCCCGTCCAAGCACCTCTGATGATGCTGGGCGATGCTGGGCGATGCTGGGCGATGCAGGATTTTTGGAGACCAGAGGTGGGGTTCCGATGCTGGGCGATGGTTGCATTTTTTTTGTATATTTTGGTGAATTGAACTATTTTTGGTTAATCAAGATGCAGATGCTAGACAAGTTGCAGATACTGGATGCCGATGCTGGACAATGTTCTTTGCTTTAATTGTATTTTTTTCCTCCACAGCTGTTGTTTTCTGCTATGTCTGCCCGAAAGACGGCTGTGGAGGAGAGGCTGGTCAAAAGGGCAACCAGGTTTCTTTGTCCATGTTTTTTTTTTTGATGAAGAAGCCTGCTCCCAGCCACTGAAGTGCTGACGCACACATGCATCTTTTTTGATTTTAACAATGTTAGAGAGATGCATGCCCATAACAGGTGCGCAACCGTCGGCAAGAGATGGAGAAGAAGCGGCAGCAGCGCAATCCCGCTTTGTCAATCACTTTTGATCCTGAAGGGGATGACCCAATGCCGGGGTCTCGGAAGGGGAAGAGTGCATTCGGGCCGAATGCCCCCCGACCATGGTTTTTCCTGAGCAAGATTTTCCCAGCAGGACAAATTGTGGCTCGAATGCCTTACTGCCACCGCTTGCCTGGACGCCTCCGACCATGGTTTTTTTGCTGTATCTGGTTCACCTGCACTTTTGTGGAGACGCTAGGGGTTTCAGAGACCTCCTTCCATTGTCAGGGGAGGGGTTTCAGAGACCTCCTTCCATTGTTAGGGCATTGTTAGGGGAGGGGTTTCAGAGACCTCCCCGATTCACAAAAAGCATTCAAAGGTGGTCAATGTGGACAAAAAGCAGCAAAAGTGATCAAAGTGGTCAAAAAGCAACCAAAAAAAAAGATGGCCAAAAGTAACCAAAAGTGGGCTAGAGTGGCAAAGGCAGCAGGCATTATAGGTACACCTTCAAGGGTTCCAAAGGCGATGCTGGGCGATGCTGGGCGATGCTGGGCCCAAGGGCATGGCTGACCAGAGGTGCTTGGTCCCGTCTGACGCATGGCTGGGCTGGTGGGTAGAGGTGGAGAGATGTATGCGCAAATGGCATTTTGTTGGCAGGGCTCCTCTCCTCTGGCACCTGCATCGGGCCCCAGGGTTTGGTTGCTTGCCCGTCAACCGTTGGGTTTGAAGGGGAGGGGTATTCTCCGAACGGGAGCCAGCCCGCTTGGTGGCGTTTCGCATGATGACCCCCTTCTCAGCGGGTTCCACTGATTAGTGATCGAGAGTGACAACCGAGCGACTGAGAGCGATGGCTGGTGATGTGTGTTCGTGCTCCATGGGCAGCGTTGGGTGGGAGGCCTGGATGGTGTTACATGATGCGCTGATTGTTCAATTCCTTTACCTGCACCGTGATGAACATTCCCAAGCTGATGGGTGCTGGGTGGTTGGAAGCACTGCGGAAACACACCACTCGTGTGGCAGAGGCATTTGGTAACCTGGGATGCGAGGCCTTGAGTGATGATTCTTCTCAGCGGGTTTCTGAGACGATGCTGAGGCTCCATAATTCTGGCTGATAAGGGCAGGAAGCAACTGGTGATTGGGGTGTGTCCAGCAGCAGGGGTTCGGAGGGTGGTGGGTGCGGTAAGTTGCAATTGTACATTCACTGAGCCTCTGTGCATCCCCTCGGTGATTTGTGTTGTGGGTGTTATGGGCAGGCAGGGGGTTTGGCATTTCGACCCCCCCCCCCGCAGGGGAGGGATCGTGTTTTGGGGTGGGCAGTGGTATAGCACACCGTCGCTGCTGAAACAGGCCGGGTCCCTAAAGTGGCCACTTCAAAAGGGAGAATTGGCTCAGTCGGAAGGCACCAGTGCCCAAGGCCGAGCCCCGCACCCAGGTAGGCACCCCATAGCAGAAGGCACCCCCCCGTCCTCCCCGGGGGCTGGGGGGTGGGGGGCTGTCGTGTGGTTGGTGGGTTATCCAATGAAAGTGCATGGGCCCCAATGCCCATCTTTGCAACCATCCACCGTTGTCTTTGCCAGTCCCGTCTGACCCGCATGGCTGGGCTGGTGGGTAGAGGTGGAGAGATGTATGCGCAAATGGAATTTTGTTGGCAGGGCTCCTCTCCTCTGGCACCTGCATCGGGCCCCAGGGTTTGGTTGCTTGCCCGTCAACCGTTGGGTTTGAAGGGGAGGGGTATTCTCCGAACGGGAACCAGCCCGCTTGGTGGCGTTTCGCATGATGACCCCCTTCTCAGCGGGTTCCACTGATTAGTGATCGAGAGTGACAACCGAGCGACTGAGAGCGATGGCTGGTGATGAGTGCTCGTGCTCCATGGGCAGCGTTGGGTGGGAGGCCTGGATGGTGTCACATGATGTGCTGATTGTTCAATTCCTTTACCTGCACCGTGATGAACATTCCCAAGCTGATGGGTGCTGGGTGGTTGGAAGCACTGCGGAAACACACCACTCGTGTGGCAGAGGCACTTGGTAACCTGGGATGCGAGGCCTTGAGTGATGATTCTTCTCAGCGGGTTTCTGAGACGATGCTGAGGCTCCATAATTCTGGCTGATAAGGGCAGGAAGCAACTGGTGATTGGGGTGTGTCCAGCAGCAGGGGTTCGGAGGGTGGTGGGTGCGGTAAGTTGCAACTGTACATTCACTGAACCTCTGTGCATCCCCTCGGTGGTTTGTGTTGTGGGTGTTATGGGCAGGCAGGGGTTTGGCATTTCGACCCCCCCCCCCCCCGCAGGGGAGGGATCGTGTTTTGGGGTGGGCAGTGGTATAGCACACCGTCGCTGCTGAAACAGGCCGGGTCCCTAAAGTGGCCACTTCAAAAGGGAGAATTGGCTCAGTCGGAAGGCACCAGTGCCCAAGGCCGAGCCCCGCACCCGGGTAGGCACCCCATAGCAGAAGGCACCCCCCCGTCCTCCCCGGGGGCTGGGGGGTGGGGGGCTGTCGTGTGGTTGGTGGGTTATCCAATGAAAGTGCATGGGCCCCAATGCCCATCTTTGCAACCATCCACCGTTGTCTTTGCCAGCCCCGTCTGACCCGCATGGCTGGGCTGGTGGGTAGAGGTGGAGAGATGTATGCGCAAATGGCATTTTGTTGGCAGGGCTCCTCTCCTCTGGCACCTGCATCGGGCCCCAGGGTTTGGTTGCTTGCCCGTCAACCGTTGGGTTTGAAGGGGAGGGGTATTCTCCGAACGGGAGCCAGCCCGCTTGGTGGCGTTTCGCATGATGACCCCCTTCTCAGCGGGTTCCACTGATTAGTGATCGAGAGTGACAACCGAGCGACTGAGAGCGATGGCTGGTGATGTGTGCTCGTGCTCCATGGGCAGCGTTGGGTGGGAGGCCTGGATGGTGTTACATGATGTGCTGATTGTTCAATTCCTTTACCTGCACCGTGATGAACATTCCCAAGCTGATGGGTGCTGGGTGGTTGGAAGCACTGCGGAAACACACCACTCGTGTGGCAGAGGCACTTGGTAACCTGGGATGCGAGGCCTTGAGTGATGATTCTTCTCAGCGGGTTTCTGAGACGATGCTGAGGCTCCATAATTCTGGCTGATAAGGGCAGGAAGCAACTGGTGATTGGGGTGTGTCCAGCAGCAGGGGTTCGGAGGGTGGTGGGTGCGGTAAGTTGCAACTGTACATTCACTGAGCCTCTGTGCATCCCCTCGGTGGTTTGTGTTGTGGGTGTTATGGGCAGGCAGGGGGTTTGGCATTTCGACCCCCCCCCCACAGGGGAGGGATCGTGTTTTGGGGTGGGCAGTGGTATAGCACACCGTCGCTGCTGAAACAGGCCGGGTCCCTAAAGTGGCCACTTCAAAAGGGAGAATTGGCTCAGTCGGAAGGCACCAGTGCCCAAGGCCGAGCCCCGCGCACGGGTAGGCACCCCATAGCAGAAGGCACCCCCGTCCTCCCCGGGGGCTGGGAGGTGGGGGGCTGGCGTGACCAAGTCGGAGGGTTGTGTGGGTGGGTTATCCAATGAAAGTGCATGGGCCCCAATGCCCATCTTTGCAACCATCCACCGTTGTCTTTGCCAGCCCCGTCTGACCCGCATGGCTGGGCTGGTGGGTAGAGGTGGAGAGATGTATGCGCAAATGGCATTTTGTTGGCAGGTCTCCTCTCCTCTGGCACCTGCATCGGGCCCCAGGGTTTGGTTGCTTGCCCGTCAACCGTTGGGTTTGAAGGGGAGGGGTATTCTCCGAACGGGAGCCAGCCCGCTTGGTGGCGTTTCGCATGATGACCCCCTTCTCAGCGGGTTCCACTGATTAGTGATCGAGAGTGACAACCGAGCGACTGAGAGCGATGGCTGGTGATGTGTGTTCGTGCTCCATGGGCAGCGTTGGGTGGGAGGCCTGGATGGTGTTACATGATGCGCTGATTGTTCAATTCCTTTACCTGCACCGTGATGAACATTCCCAAGCTGATGGGTGCTGGGTGGTTGGAAGCACTGCGGAAACACACCACTCGTGTGGCAGAGGCATTTGGTAACCTGGGATGCGAGGCCTTGAGTGATGATTCTTCTCAGCGGGTTTCTGAGACGATGCTGAGGCTCCATAATTCTGGCTGATAAGGGCAGGAAGCAACTGGTGATTGGGGTGTGTCCAGCAGCAGGGGTTCGGAGGGTGGTGGGTGCGGTAAGTTGCAATTGTAAATTCACTGAGCCTCTGTGCATCCCCTCGGTGATTTGTGTTGTGGGTGTTATGGGCAGGCAGGGGGTTTGGCATTTCGACCCCCCCCCCCGCAGGGGAGGGATCGTGTTTTGGGGTGGGCAGTGGTATAGCACACCGTCGCTGCTGAAACAGGCCGGGTCCCTAAAGTGGCCACTTCAAAAGGGAGAATTGGCTCAGTCGGAAGGCACCAGTGCCCAAGGCCGAGCCCCGCACCCAGGTAGGCACCCCATAGCAGAAGGCACCCCCCCGTCCTCCCCGGGGGCTGGGGGGTGGGGGGCTGGCGTGTGGTTGGTGGGTTATCCAATGAAAGTGCATGGGCCCCAATGCCCATCTTTGCAACCATCCACCGTTGTCTTTGCCAGTCCCGTCTGACCCGCATGGCTGGGCTGGTGGGTAGAGGTGGAGAGATGTATGCGCAAATGGAATTTTGTTGGCAGGGCTCCTCTCCTCTGGCACCTGCATCGGGCCCCAGGGTTTGGTTGCTTGCCCGTCAACCGTTGGGTTTGAAGGGGAGGGGTATTCTCCGAACGGGAACCAGCCCGCTTGGTGGCGTTTCGCATGATGACCCCCTTCTCAGCGGGTTCCACTGATTAGTGATCGAGAGTGACAACCGAGCGACTGAGAGCGATGGCTGGTGATGAGTGCTCGTGCTCCATGGGCAGCGTTGGGTGGGAGGCCTGGATGGTGTCACATGATGTGCTGATTGTTCAATTCCTTTACCTGCACCGTGATGAACATTCCCAAGCTGATGGGTGCTGGGTGGTTGGAAGCACTGCGGAAACACACCACTCGTGTGGCAGAGGCACTTGGTAACCTGGGATGCGAGGCCTTGAGTGATGATTCTTCTCAGCGGGTTTCTGAGACGATGCTGAGGCTCCATAATTCTGGCTGATAAGGGCAGGAAGCAACTGGTGATTGGGGTGTGTCCAGCAGCAGGGGTTCGGAGGGTGGTGGGTGCGGTAAGTTGCAACTGTACATTCACTGAACCTCTGTGCATCCCCTCGGTGGTTTGTGTTGTGGGTGTTATGGGCAGGCAGGGGTTTGGCATTTCGACCCCCCCCCCCCGCAGGGGAGGGATCGTGTTTTGGGGTGGGCAGTGGTATAGCACACCGTCGCTGCTGAAACAGGCCGGGTCCCTAAAGTGGCCACTTCAAAAGGGAGAATTGGCTCAGTCGGAAGGCACCAGTGCCCAAGGCCGAGCCCCGCACCCGGGTAGGCACCCCATAGCAGAAGGCACCCCCCCGTCCTCCCCGGGGGCTGGGGGGTGGGGGGCTGTCGTGTGGTTGGTGGGTTATCCAATGAAAGTGCATGGGCCCCAATGCCCATCTTTGCAACCATCCACCGTTGTCTTTGCCAGCCCCGTCTGACCCGCATGGCTGGGCTGGTGGGTAGAGGTGGAGAGATGTATGCGCAAATGGCATTTTGTTGGCAGGGCTCCTCTCCTCTGGCACCTGCATCGGGCCCCAGGGTTTGGTTGCTTGCCCGTCAACCGTTGGGTTTGAAGGGGAGGGGTATTCTCCGAACGGGAGCCAGCCCGCTTGGTGGCGTTTCGCATGATGACCCCCTTCTCAGCGGGTTCCACTGATTAGTGATCGAGAGTGACAACCGAGCGACTGAGAGCGATGGCTGGTGATGTGTGCTCGTGCTCCATGGGCAGCGTTGGGTGGGAGGCCTGGATGGTGTTACATGATGTGCTGATTGTTCAATTCCTTTACCTGCACCGTGATGAACATTCCCAAGCTGATGGGTGCTGGGTGGTTGGAAGCACTGCGGAAACACACCACTCGTGTGGCAGAGGCACTTGGTAACCTGGGATGCGAGGCCTTGAGTGATGATTCTTCTCAGCGGGTTTCTGAGACGATGCTGAGGCTCCATAATTCTGGCTGATAAGGGCAGGAAGCAACTGGTGATTGGGGTGTGTCCAGCAGCAGGGGTTCGGAGGGTGGTGGGTGCGGTAAGTTGCAATTGTAAATTCACTGAGCCTCTGTGCATCCCCTCGGTGATTTGTGTTGTGGGTGTTATGGGCAGGCAGGGGGTTTGGCATTTCGACCCCCCCCCCCCGCAGGGGAGGGATCGTGTTTTGGGGTGGGCAGTGGTATAGCACACCGTCGCTGCTGAAACAGGCCGGGTCCCTAAAGTGGCCACTTCAAAAGGGAGAATTGGCTCAGTCGGAAGGCACCAGTGCCCAAGGCCGAGCCCCGCACCCAGGTAGGCACCCCATAGCAGAAGGCACCCCCCCGTCCTCCCCGGGGGCTGGGGGGTGGGGGGCTGGCGTGTGGTTGGTGGGTTATCCAATGAAAGTGCATGGGCCCCAATGCCCATCTTTGCAACCATCCACCGTTGTCTTTGCCAGTCCCGTCTGACCCGCATGGCTGGGCTGGTGGGTAGAGGTGGAGAGATGTATGCGCAAATGGAATTTTGTTGGCAGGGCTCCTCTCCTCTGGCACCTGCATCGGGCCCCAGGGTTTGGTTGCTTGCCCGTCAACCGTTGGGTTTGAAGGGGAGGGGTATTCTCCGAACGGGAACCAGCCCGCTTGGTGGCGTTTCGCATGATGACCCCCTTCTCAGCGGGTTCCACTGATTAGTGATCGAGAGTGACAACCGAGCGACTGAGAGCGATGGCTGGTGATGAGTGCTCGTGCTCCATGGGCAGCGTTGGGTGGGAGGCCTGGATGGTGTCACATGATGTGCTGATTGTTCAATTCCTTTACCTGCACCGTGATGAACATTCCCAAGCTGATGGGTGCTGGGTGGTTGGAAGCACTGCGGAAACACACCACTCGTGTGGCAGAGGCACTTGGTAACCTGGGATGCGAGGCCTTGAGTGATGATTCTTCTCAGCGGGTTTCTGAGACGATGCTGAGGCTCCATAATTCTGGCTGATAAGGGCAGGAAGCAACTGGTGATTGGGGTGTGTCCAGCAGCAGGGGTTCGGAGGGTGGTGGGTGCGGTAAGTTGCAACTGTACATTCACTGAACCTCTGTGCATCCCCTCGGTGGTTTGTGTTGTGGGTGTTATGGGCAGGCAGGGGTTTGGCATTTCGACCCCCCCCCCCCCCGCAGGGGAGGGATCGTGTTTTGGGGTGGGCAGTGGTATAGCACACCGTCGCTGCTGAAACAGGCCGGGTCCCTAAAGTGGCCACTTCAAAAGGGAGAATTGGCTCAGTCGGAAGGCACCAGTGCCCAAGGCCGAGCCCCGCACCCGGGTAGGCACCCCATAGCAGAAGGCACCCCCCCGTCCTCCCCGGGGGCTGGGGGGTGGGGGGCTGTCGTGTGGTTGGTGGGTTATCCAATGAAAGTGCATGGGCCCCAATGCCCATCTTTGCAACCATCCACCGTTGTCTTTGCCAGCCCCGTCTGACCCGCATGGCTGGGCTGGTGGGTAGAGGTGGAGAGATGTATGCGCAAATGGCATTTTGTTGGCAGGGCTCCTCTCCTCTGGCACCTGCATCGGGCCCCAGGGTTTGGTTGCTTGCCCGTCAACCGTTGGGTTTGAAGGGGAGGGGTATTCTCCGAACGGGAACCAGCCCCCTTGGTGGCGTTTCGCATGATGACCCCCTTCTCAGCGGGTTCCACTGATTAGTGATCGAGAGTGACAACCGAGCGACTGAGAGCGATGGCTGGTGATGTGTGCTCGTGCTCCATGGGCAGCGTTGGGTGGGAGGCCTGGATGGTGTCACATGATGTACTGATTGTTCAATTCCTTTACCTGCACCGTGATGAACATTCCCAAGCTGATGGGTGCTGGGTGGTTGGAAGCACTGCGGAAACACACCACTCGTGTGGCAGAGGCACTTGGTAACCTGGGATGCGAGGCCTTGAGTGATGATTCTTCTCAGCGGGTTTCTGAGACGATGCTGAGGCTCCATAATTCTGGCTGATAAGGGCAGGAAGCAACTGGTGATTGGGGTGTGTCCAGCAGCAGGGGTTCGGAGGGTGGTGGGTGCGGTAAGTTGCAACTGTACATTCACTGAGCCTCTGTGCATCCCCTCGGTGGTTTGTGTTGTGGGTGTTATGGGCAGGCAGGGGGTTTGGCATTTCGACCCCCCCCCCCCGCAGGGGAGGGATCGTGTTTTGGGGTGGGCAGTGGTATAGCACACCGTCGCTGCTGAAACAGGCCGGGTCCCAAAAGTGGCCACTTCAAAAGGGAGAATTGGCTCAGTCGGAAGGCACCAGTGCCCAAGGCCGAGCCCCGCGCACGGGTAGGCACCCCATAGCAGAAGGCACCCCCGTCCTCCCCGGGGCCTGGGGGGTGGGGGGCTGGCGTGACCAAGTCGGAGGGTTGTGTGGGTGGGTTATCCAATGAAAGTGCATGGGCCCCAATGCCCATCTTTGCAACCATCCACCGTTGTCTTTGCCAGCCCCGTCTGACCCGCATGGCTGGGCTGGTGGGTAGAGGCGGAAAGATGTATGCGCAAGTGGCATTTTGTTGGCAGGTCTCCTCTCCTCTGGCACCTGCATCGGGCCCCAGGGTTTGGTTGCTTGCCCGTCAACCGTTGGGTTTGAAGGGGAGGGGTATTCTCCGAACGGGAACCAGCCCGCTTGGTGGCGTTTCGCATGATGACCCCCTTCTCAGCGGGTTCCACTGATTAGTGATCGAGAGTGACAACCGAGCGACTGAGAGCGATGGCTGGTGATGTGTGCTCGTGCTCCATGGGCAGCGTTGGGTGGGAGGCCTGGATGGTGTTACATGATGCGCTGATTGTTCAATTCCTTTACCTGCACCGTGATGAACATTCCCAAGCTGATGGGTGCTGGGTGGTTGGAAGCACTGCGGAAACACACCACTCGTGTGGCAGAGGCACTTGGTAACCTGGGATGCGAGGCCTTGAGTGATGATTCTTCTCAGCGGGTTTCTGAGACGATGCTGAGGCTCCATAATTCTGGCTGATCCGGGCGGCAGGCAACTGGTGATTGGGGTGTGTCCAGCAGCAGGGGTTCGGAGGGTGGTGGGTGCGGTAAAGTTTGGTCTTTTTTGGGTGGGTAGGGACCCGGGTACCCACCAACTTTTTTTTGACCCCCTGTCCAACGACCACTACCCCCAGTCAGCGGACAGAGGTTACCGGTCATTGGACATATTGGGGGGGGTAACAAAAAAAAGCGAAAAAGGTGAAAGATACAAGAGAAATTACTGCATGGAATTGCTTCTGCGCAACTGTGGCCAACGAAGGGTGCATCCAGGCCCAGGCAAGTTGTTTGACTGGTTTTTTCTTTTGTACCGTCTGGTTTTGCTCCACACTCTTATCCCCGGGATGGTGGTCAAACACCTATGAATTGGCTGGGCGTGACCGTGTGTGAACAATTTTGCAGAACCCAAAGGCGGGACTTCATGACTGGGCGTGACTTAGCATGTCTGGTTTATTTTTGACATTTTTGTGGTTTTTGGCTGTTTATGGGCCTTTTAACAATATATGACGTCCTTTTTTTGAACAATTTTGACAGCTTTTTGCTATTTTTGAAATTAACATCAGCAAATATTGGCCAAAAACAGCCAATAAAAGCCACAAGTTTGCAAAAATGGCCAAGAAACCCATGCCTGAGTTGGTATGACTCAGAATGTTGTTGGCGTGTTTGGGCGTGACTGTGACGTTTGGGGTGGCCGAAAGTGTCACTTGCCGCCGAAGGCAGTGGAGCTTTTTGGGCGTTTTTGACCAAAGTCCATTATTTCCGTCCATTTTGGACGGAGTGGCCAAAGTGACTAACACCCCACAGGGGGTATGATTGGTATTTTGAAAGTGGAATGCGAATCCCCCCGGTTTGGTCCACCCCCGGCCGGGGGGTTTAGGACCATTTGGGCCGAAAATTTCTCTCTTCCCAGCACTTGGCAGTTTTATGTATTATCGCCTGTTGTTGTCGATCCGTTGGACTTAAAAACGCAACCTTGCGGCATTTTTTTGAAGCCCCTCAAAAACGCTTTGACGATTTGCGCCAGCATTTTTCGCGACTTTTTGCCCCTTTTTGGGGGAAAATTTTTTGGGTGGGTCCCCTGGGACCTCCCCTCCTCCCTGGTTTAAAAAAACCCCTGAACCCCCCTCTGCTCCCGCCGGGCTGTACGGGAAACCTCATTGGTGGTGGGTTGGGGAGGTCTCCGACCCCCCCCCCCATCCTGGTGGAGGGGTTCGACCATCTCCCCACCATCCCTTCCCGTGTTTCCAGGGGCACCAATTTTTAGGTGAAAGTTTTCTTTCGGTGGCGAAGAAAAGTTTTTCCCTTGGGCAGCACCATGGTGGGTGGGGGGGGGTCGACCCCCCCCCCCCCACCCGGCCTTCCCGTACCTCCCTGTCCGGCGAACAATAAGGGGGGGTGAAAAGGGGAAAGCCAAGGCGCAGCCCAAAAGCTTTCCGTCTGCCGGGGCCCAGGTGCAAGAAGGTGGGGGACCATGAGGTGGGTATTTCCCTTCCGGAATAACCCCTGCTTCCCCTTTTCGTAGGAGAGGATCCAGAACCTCACAGTTCGAGAAGTCCGGCAGGCCATCCCTGCGCACTGCTTCAAGCGGTGCACCTACCTCTCAATGATGCACTTCGCCTACGACGTCTGTGTGATGTTGGCCACTGCCGCCGCCGTTATTTTTGCCTGGCGGAACCTCAGTGGTTGGTGGATGGCTATCATTTGGCCAGCATATTGGTGGTATCAGGTTTGTTCCTCTCGCTGGAATATTCCCTCAATCGAGAATGTGTGCGGCATGTGGGGATATTGGGTTTTGTTTATTCCAAAAAATTGATTTTTGTGGTTTTGATAATGGATTAATGGATAATGAATTTTGTTTTCCGGTGACCTGGAGGGTGGGTTCTGGCCTGCCCCGGAAAAACAGCCCAAATTGGAATTGTGACCCACTTTCCCACAGAAGTCCCCCTTTTGGTTTAACAAAAGTTTGCTGAGGGGGGTTGCCAACCAGGAACTCCCCCCTCCAGTCAGCATCCCTGGAAATCCGAGATGGACCAAGGGGAAACAAGGCCCAAGTTGTGGAATGTCTACCAAAAACACTGGGTTGTCAGAATCCAGGGCAGTGATGTGCAGGCTACAGTCTCGTGCAAATTTTGTCACTTGAGAACATCAGCTAGTCAAACTCATGGTCTTTTCAGGGCTTGAATGGGACGGCCCTCTGGGTCTTATCCAGGCTTCTGTTTGAGGAAGGTTGCACCCGGGTGGGGGGAGTTTTAGGATGGGTGGTCTTATGGTGCAACTCCCAGCACCAAAGCAGAGATTTTTTTCAGTTTCGTGCCCCGGGGGTCCAGATTTTCATTTATTGCGCCGCGGAAATTTTTTTTCTGCGCCGCGTGCACCGCGGCACCGGAAATTTCTTTCTGTTTTTTTTGAGATTTTCTGAGATTTCTTTCTTTCTCTGCGCCGCGGCAGCAGCGCGTGCACACTGCATGTGTCGCGGGAACGTCATTGGTGCGCCGCGGAATTTTTTTTTTGCGCCGCCGCAAACTTCGCCACCTCCACCTCCACGAGTGATGTTGCCACCTCCACGTACACAGACAAAAATGGCCAAAGGCAAAAAAACAGCGCCAGGCCAGAAGAAGGGATAAAAGGCACCAAACAAACAGCCACACTGCACACAGGGGGGGTGGACTAGTTTTTTCGGAGTGACTGACTAGTTTTCGGCTCCAGACCCCCCCCCCTGGGGGGTGGGGTTGGCCAGGATTGGCCGAAAGTGGCCACCCGGCAGCGAAGGTGCTAATTTTTTTGGGGGGTTTTGGTCAAAATGCATCGATTTTGAGCCATTTTTGGCTGAGTGGCCAAAGTGACCGAACCCCCGGGGGGGGGGGGTAGGAAGGGCGACTTTTTTGAAGAGAGATAGAGTGGTTTTTTTAAAAGTGCCCAAAATGTGATTTTTGACTGTCAATTTTTTTCGGCCCGGCGCTTCGCGCCAGGTCGGTCACCCAGACCGCGCCATTGTTTAGGTGTTGGGGGGTGCCCCCCCCCCCTCCTGTACCGGGGGGCCTGGACTCCCCCCCCCCCCCTACATCGAAACCTGGCCTGGACTTCCAGCACCGAAGCACGTGGGCCTGATAACGGATATTGTAAAGGAAAGTTTGCCTGGGTCGGCAGACATTTGCTCCCAAAACCGACCCGGGTGCTTTTAATGTGCAGGCACTCCAAAGCACAACCCCCCTGGGTGGTAAAAAGGAAGCCCGGTCTTGGCCCATGAATGTGGTCACGGCGGCTTCACCGATTCCAAGCTTATTAATGACATCGTGGGCTTCATCACCCACTCTTTCCTCCTAACCCCCTACTGGTCCTGGGCCCTCATCATACCGTTGCTGCCCAAGGGGGTGACTGACTGGTCTCGGTGCGACGGGTTTTTTGTTTTTTTGGGCTCCCGATCTGGGCTGTACATAAATCCCGGAAGCCCCATAGGTGGTGGTGGGGTCCAACTGGGTGTGCCGGGTGGTGGTGGGGGTCGACCACCTGCCTGCACAGCCTTGCCCAGGACCCAGGCACCAAAGGCATTGGTATTTTTTTGAGAAAATTCAGGGGGGGAAATGCGTTTCGCAATTTTTCGCATTTTTTTCGCATTTTTTGCGCATTTTTCAAATTTTCCAAAAATGCGCATTTTTTCGCATTTTTTGCGCATTTTTTTGCTTTTTTTTTTCAGCGCACTTTTTTTGCTTTTTGGCAAAATTTTTCGCATTTTTTGACCATTTTTGCAATTTTTTTGCAAAAATTTGAAATTTTCGACTTTTTTTTGAAACTTATATTTTCCTCATTATTCTACATAATTTCTTCAAATTTGGTGCCATTTTTGCATGTTTTTGGGTAATTTTTGCATTTTCGTGATTTTTGCGCATTTTTTCTCGGATTTTGCACATTTTTGCGCATTTTTTCCAAAAATGCGCATTTTTTGCGATTTTTTGCGCATTTTTTTCCGAGGCCGGGCGAGCGCATTTCCCCCCCCCCCCCGGAAAATGCAATGATTGTTTTGCTTGACTGACCACCCAGGGCGGGCAGTGGGTGGTTTGGTTTTCAAGAAGAGACCGACCGTCTGGTTTTTTTGTTACCTGCCCCCCCCCCCCCCTGTCCACCGACCACCCCTATGTACCGACCCCCCCCCCCCCCGGTCGGGGGTTGCAGGTTGGGGGGGTATACGACCCCCCCCCGGTTGGGGGTTACAGGTGACCACGAAAGTTTTTTTTACCTCTCATGTGCTTGTCGACCAGCTTTTTCCCGTGTTTCCTGTCCATTATATACTGAGAGGCGCCCGTGGCAACCTCCACCGTTATGAAATTGCCTTTTTTTGGCTTTTGGTTTTGTTTTGGTCCCTAGTGGTACCATGATGACCACCCCCCGGTCGGGGTTGCAGGAACCCGGTCGTTGGACGGGGGGGGGGGGCGGGTAATGGAAAGTGAGTCCTGGGGGTCGATGAGAAACGGTATGGCCCTGACCCACGCCAAGCACCACCGCCGCACCAACCACATTTCTGAGGGGGAGACCTGGGTGCCTGCCATCACCCGCAACCCCGACAAGCCAAAGATCAAGTTCTTCAAAACCCACCTCGGGACTTGCATCCGCATTACCATCGTCTGGACCGTTGGGTGGTACCTGTATTTGTTCAGGAACGACACAGGTGTGAGGCCTCTGTTTATTGTTTTGCGGTTGATGTTTTCTGTCAGTTACCCAACCAGTGTCCGCGCTGGCAGCCATTCTTTTTGATTCTGCGCTCTGGATGGAGGCGTCCTCATCCAGACCACTGTTTTAATTGAAAATATGACCAGGGGTCATCTATTTAATCTGTCCAGTCGATATTTTTTTTTCATTTTGGCGACAGTGGCGAGAGTGACCCCTCCCCAGGGTTCGATATAAGGAGGGCGTACATGGTTTAGGTGGGGGGGTGTCCCCCCCCACCCCCCCCCCCCTCCTGGCCCGCAGGGGGGGGTCCGGACCCCCCCCCACCTGCAAGGGGGGTCCGGACCGACCCTGGGGGGCAAAAATTGACCCCCCCAAAAAGGGTCATTTTTTTCCTCCGGCGCTTTGCGCCGAAAATTTTTTGCATTTTTGGCACATTTTTGCCCCAAAAGTGCCCAAAATGGCACTTTTGGTCGTTTTGGGGGGTCAAAAATTTTCGGCCACCCTGACCGCCGCACCTGTTTAGGTGGGGGGGGGTGTCCCCCCCACCCCCCCCCCCCCCCTCCTGGCCCGCAGGGGGGGGTCCGGACCACCCCCCCCCACCTGTCCTTATATCGAACCCTGCCCCTCCCCCCCCCCAGGGGCAGGGCTGTGCGGGAATCCCCAAAACCCTATGGGTGGTGGGCAGATTTTCATTTTTTTCCGCCGCGGAATTTTTTTTTGTGCCGCGGCAAACTTCGCCACGGCGCACTAAATGAATGCCTGGTGATGGGTTTTGGGGGTCCGACCCCCCCCCACAAGGCTTTTTGTCAAAAGTTTTCCCTCGGCGCTCACGGCGCTGAAGGAATTTCTTCTTGCACAGTGCTGTGGTGGGGACGTGGGGGGGGTGCGATTAAAAGTCAAAGTCCACGTCGAAGCAATTAAAAAAAAACGTCGTTACAGAGTGGCGTATTGGCAATCCGTTCTGGGAGTGCCGCGAGTGCATGTTGGTCACATGTGACCCATCAAAACTGCCACGTTCACCCCCCCCCCCCCCCCCGACCCGGCCCCCCCCCCCCGACCCGGCCCCCCCCCCCGCGACCCCCCCCCCCGACCCCCCCCCCCCGACTCACACACCCCCTGCATCAATTGTGTCGATCCAGGGCTGCATAGAGACCCCTTGGCCGCCGCTGTTTTCCGCAGGCAGCTTCAAGAACCACGGCCAGTCGCATTTCAACCCCAACTCCAAGGGCCTGTTCCGCCCGGCGGACCGCCCCTGGGTGCTGCTGAGCGACGTCGGGATGATCGTCACGCTGATCGGCCTGGCGGCGGCCAGGACGGCTTTGCCCCCCCCCCCCCCCCCGACGGTGGTGCCAGGCCTGCCTCGGTCAAGCCCATCCCCCACCCTGCCACCCACCCCCACCCAACAGTGAAGGGGTGGGTGCAAATGATGTTGGGGGATTTTGGGCCATTTTCCCAGTAGAGTTGCAATGTGCCTGATTTCTTTCCCAATTTTGATGCAGCGGGTCATAAATTAACGTCAGCCAAGAGGAATGCTCGGTCTGTCTTGGCAGAAATGACATTGGCTCTGGCACGGCCAACTGCCTGCCAAGGTCATTGGGCAAACAAATTTCTTGCCCTATGTTGATTTGTCCAACAAGAGCTGCCTTTTGACCTGGCTTGGCGCAGTTGCTGGAATGGCAGTGGTTTTGTCCCTCGGTTGTGTCTGCATGCATTTGTACACCTTTTGTATGCGGAAAAAGGTCGCCCTTGGAGCTGGCAGTGTTGAAGTTTGGGCTGGTGCCGGTGCTCCTGGTCTATGCCATCCCCCAGATGATCACTGATATGTACCTGGTGTCCATCACATTCATGCAGCACGCCCATCTGGACCTGCCGCATATGGATTTCCCTGTGTGGAATTGGCTGAACGGTGCACTTTGCACCGTGGACAGGTGCGTTGTGTTGTGCGGAGTGAGTTGGGGAGGTGCGATGAAATGTGGTCAGACTTTTTTTTGGGTTTTGTGACCGGGGGTCCCGCCGAGCCCCCCCCCCCGGGGGGGGGTTGGGTCTGGAATTACACCAGAGTTGCTCACCCGGCCAGGCGGAGCAACAAAAGAGCACTGGTGTAAGGGGTACCTGGCCACTCACCATACAGCAAGCTGGGCCAGAAGCGAAGTTAGCGCCGGCCATAAACAGACAAGAAGCCTTAGTGCAACTAAGGGGGTTGGGCAGATTGGGAACCAGATATTCTTTTAAGTTTTGACCCGTCAGGGTCCTGGCCGAGCCCCCCCCCCCCCCCCCCGGGGGGGTCGGGCCCAGTGGGTCCCCCACGGGGGGGGGGTCGGGGACATGCAGGGATGGATGGGAAGCCCTTCGGTATTGAGGTAGGCGGTAGGGGGGGGTCCCGACCACCCCCCCCCCCCTGGGTGGTCGAGGCATCGTTTCAGATGTGCCGCGGCAAAATTTGATTGTGTGCCACGTGCGCCGCGGAAATTTTTTTGTGCCGCGGCAAACTTCGCCGCTAAATGAAAGTCTGTTACATATGTTACGTGTTTCTTGATCTTTTTGGGAAAAACTGCAGTGGAAATAGTGTCCCTCCCCACCGACCACCTTTGCATACCAATCACCGACCTGCCACCCAGGGGGCGCAAGACCAGGCGCAAATACCCCAATAAACAGAAAATTTTTAACTCTGCTGAAGCGTTTTGCATATGACCACCCTCAAACCAGCAATCCTGGTCGGTTCTTGGCACCCTTTAATCTGCAACCTCCCGGGGGGTGACCCCCCCCCCCCCGGTTGGGGTTGTCGGAACACGGTTGGTGGGAAGCAAATTTGAATTCCCAGGGGAACAGAGTTCTTTTGTGTTGAAAATTTTCATCGACCACCGCTTGATCATGCAAAGTACCCATCTCACCCCTGTTGCAATGCTGCGCTTTGGTTTGGTGGTGGGTGTGGTGCTTCAGTGATTTGCTCCCCGCACAGGTCGATGGGCCCGTGGGTGGATTCTAAACTGCACCATATTGTGGACAGCCATGTGGTGCACCACATTTTTTCTGATATGCCATTCTATGGGGCGAAGGAGGCAACGCCTTACAACCAGGCCTCTTTATGCACAAGTTCCTCAAGACCGCTTTTTACCCATGTCAGGTACCATTGTCTTAGATCACCACATCATTTGAGTGATGGTGGAATTAACATTTGCACATCGAAAAATCCAACATTTTGGGCAAGTGTTTTTTTTATCTGAGAACAAGCTTTTTGACAATTTTTTGCCACATTCACAAATTTCTCCCCCTTCAATCCCACATTTTTTTTCAGCGATTTCAAACAAGTTTCTTGTTTGAATGAGGATGTTCCCAAATGTGAACATCGTGATTCTTTGACCATGTGGACCCTATGTGTTGTGAAAAAAAAAACAGAAATGTGCGCAATGTTTGATGAAAGAAGTCAGAAAAAACCTGCATTTTCATTTTCATTTCTGCATTTGAGCCTGCTCCTGTTGCCAAGTACCTCACCGACAAGTACGGTGAGGGGGTATACAACTCCAAGCAGACCAGCTATTTGGGGTATTGGCGGGACTTCTACCAGGCCATGAAGGAGGCAATTGTGGTGGTGCGGGGTCGTTCCATTGCTTGCGTTCATCCACATCCTTGTCCCTGTCTCACCATCATCGTGCCGGAAAGAGCTGCCCCCCCCAAAGCTGCATTTTTGCACATTTTTGGTTGGAGTCATTGTAGCCGA
