## Supplemental Table 4 for "Identification and characterization of the delta-12 fatty acid desaturase from *Euglena gracilis*"

**FAD12**

**CAAGGAGCACCTGCGGAAGT = gRNA, A is the deleted nucleotide after CRISPR knockout**

***TGG* is the PAM sequence**

ATGGCAGCCGCTGTGGGATTGGTTTGGAACCGCAAGCGCGCCACAGCGGTGTGGAAGGCGACGCAGGAACGGCGCGT**CAAGGAGCACCTGCGGAAGT*TGG***AGGAGAGCAAGCAGGCGGCGGCCAAGGCCGCGGCGGCGGAGGAGGACGACACGGACCCCCTGGACGCCCTGGACAGCCTGGACGCCCTCGGGGATAAGAAGGAAACCTCTTCCCTCCGGGCCCGCAAGGTTGCGCAGAAGGCCGCGGAGGGGGCGGAGAAGGTGGTGGAAGGGGAGAAGAAAAGCGTTCTGGAGAGGATCCAGAACGTCACAGTTCGAGAAGTCCGGCATGCCATCCCCGCGCACTGCTTCAAGCGGTGCACCTACCTCTCAATGATGCATTTCGCCTACGACGTCTGTATGATGTTGGCCACTGCCGCCGCCGTTATTTTTGCCTGGCGGAACCTCAGTGGTTGGTGGATGGCTATCATTTGGCCAGCATATTGGTGGTATCAGGGCTTGAATGGGACGGCCCTCTGGGTCTTGGCCCATGAATGTGGTCACGGCGGCTTCACCGATTCCAAGCTTATTAATGACATCGTGGGCTTCATCACCCACTCTTTCCTCCTAACCCCCTACTGGTCCTGGGCCCTGACCCACGCCAAGCACCACCGCCGCACAAACCACATTTCTGAGGGGGAGACCTGGGTGCCTGCCATCACCCGCAACCCCGACAGGCCAAAGGTCAAGTTCTTCAAAACCCACCTCGGGACTTGCATCCGCATTTCCATCGTCTGGACCATTGGGTGGTACCTGTATTTGTTCAGGAACGACACAGGCAGCTTCAAGAACCACGGCCAGTCGCACTTCAACCCCAACTCCAAGGGCCTGTTCCGCCCGGCGGACCGCCCCTGGGTGCTGCTGAGCGACGTTGGGATGATCGTCACACTGATCGGCCTGGCGGCGGCAGTGTTGAAATTTGGGCTGGTGCCGGTGCTCCTGGTCTATGCCATCCCCCAGATGATCACTGATATGTATCTGGTGTCCATCACGTTCATGCAGCACACCCATCTGGACCTGCCACATATGGATTTCCCTGTGTGGACTTGGCTGAACGGTGCACTTTGCACCGTGGATAGGTCGATGGGCCCGTGGGTGGATGCCAAACTGCACCATATTGTGGACACCCATGTGGTGCACCACATTTTTGCTGATATGCCATTCTATGGGGCGAAGGAGGCAACGCCCTACGTTGCCAAGTACCTCACCGACAAGTACGGTGAGGGGGTATACAACTCCAAGCAGACCAGCTATTTGGGGTATTGGCGGGACTTCTACCAGGTCATGAAGGAGGCAATTGTGGTGGTGCGGGGTCGTTCCATTGCTTGCGTTCATCCACATCCTTGTCCCTGTCTCACCATCATCGTGCCGGAAACAGCTGCCCCCCCCCCCAAAGCTGCATTCTTGCACATTTTTGGTTGGAGTCATTGTAGCCGATGA

EG FAD12 knock out CRISPR translation showing premature stop codon

*** = Stop codon**

MAAAVGLVWNRKRATAVWKATQERRVKEHLRSWRRASRRRPRPRRRRRTTRTPWTPWTAWTPSGIRRKPLPSGPARLRRRPRRGRRRWWKGRRKAFWRGSRTSQFEKSGMPSPRTASSGAPTSQ*****CISPTTSV*****CWPLPPPLFLPGGTSVVGGWLSFGQHIGGIRA*****MGRPSGSWPMNVVTAASPIPSLLMTSWASSPTLSS*PPTGPGP*****PTPSTTAAQTTFLRGRPGCLPSPATPTGQRSSSSKPTSGLASAFPSSGPLGGTCICSGTTQAASRTTASRTSTPTPRACSARRTAPGCC*****ATLG*****SSH*****SAWRRQC*****NLGWCRCSWSMPSPR*****SLICIWCPSRSCSTPIWTCHIWISLCGLG*TVHFAPWIGRWARGWMPNCTILWTPMWCTTFLLICHSMGRRRQRPTLPSTSPTSTVRGYTTPSRPAIWGIGGTSTRS*RRQLWWCGVVPLLAFIHILVPVSPSSCRKQLPPPPKLHSCTFLVGVIVAD
